# A haplotype-resolved genome assembly of the Nile rat facilitates exploration of the genetic basis of diabetes

**DOI:** 10.1101/2021.12.08.471837

**Authors:** H. Toh, C. Yang, G. Formenti, K. Raja, L. Yan, A. Tracey, W. Chow, K. Howe, L.A. Bergeron, G. Zhang, B. Haase, J. Mountcastle, O. Fedrigo, J. Fogg, B. Kirilenko, C. Munegowda, M. Hiller, A. Jain, D. Kihara, A. Rhie, A.M. Phillippy, S. Swanson, P. Jiang, D.O. Clegg, E.D. Jarvis, J.A. Thomson, R. Stewart, M.J.P. Chaisson, Y.V. Bukhman

## Abstract

The Nile rat (*Avicanthis niloticus*) is an important animal model for biomedical research, including the study of diurnal rhythms and type 2 diabetes. Here, we report a 2.5 Gb, chromosome-level reference genome assembly with fully resolved parental haplotypes, generated with the Vertebrate Genomes Project (VGP). The assembly is highly contiguous, with contig N50 of 11.1 Mb, scaffold N50 of 83 Mb, and 95.2% of the sequence assigned to chromosomes. We used a novel workflow to identify 3,613 segmental duplications and quantify duplicated genes. Comparative analyses revealed unique genomic features of the Nile rat, including those that affect genes associated with type 2 diabetes and metabolic dysfunctions. These include 14 genes that are heterozygous in the Nile rat or highly diverged from the house mouse. Our findings reflect the exceptional level of genomic detail present in this assembly, which will greatly expand the potential of the Nile rat as a model organism for genetic studies.

## Introduction

Model organisms are essential tools for the mechanistic understanding of human physical and mental health. The high-quality genomes of house mouse (*Mus musculus*) (M. G. S. Consortium and Mouse Genome Sequencing Consortium 2002) and Norway rat (*Rattus norvegicus*) (R. G. S. P. Consortium and Rat Genome Sequencing Project Consortium 2004) have enabled researchers to discover important molecular mechanisms in biological processes that have been applicable to human health. However, not all traits are well modeled by these organisms, making additional model organisms necessary for translational studies. A diurnal rodent, the Nile rat, also known as the Nile grass rat or African grass rat, is a promising model organism to address the translational gap between animal research and human biology, particularly in two areas – circadian rhythms and type 2 diabetes.

Both house mouse and Norway rat are nocturnal, while humans are diurnal. The difference between the two chronotypes is more complex than a simple flip in daily activity pattern, and likely involves distinct wiring of neural circuit and gene-regulatory networks (Yan, Smale, and Nunez 2020; Kalsbeek et al. 2008; Langel et al. 2018). Nile rats naturally exhibit clear diurnal patterns in behavior and physiology (Yan, Smale, and Nunez 2020), and have retinal anatomy as well as large retinorecipient areas in the brain typical for animals active during the daytime (Gaillard, Karten, and Sauvé 2013; Gaillard et al. 2008). This makes the Nile rat an important model organism for metabolic, cardiovascular, inflammatory, neurological, and psychiatric disorders in which circadian disruption is a risk factor and/or a hallmark symptom (Cederroth et al. 2019).

Additionally, the Nile rat has been developed as a model of type 2 diabetes. Nile rats live without diabetes on their native diet comprising mainly leaves and stems (Sibly and Brown 2007) or laboratory high fiber diets (Bolsinger et al. 2017; Toh, Thomson, and Jiang 2020). However, they rapidly develop diet-induced diabetes when fed on a conventional energy-rich laboratory rodent chow (Chaabo et al. 2010). Conversely, house mouse and Norway rat are relatively resistant to diet-induced diabetes requiring extreme high fat diets, chemical intervention, or genetic manipulation to develop diet-induced diabetes (Kottaisamy et al. 2021). Diabetic Nile rats also recapitulate the natural progression of type 2 diabetes in humans (K. Yang et al. 2016) and reproduce clinical retinopathy features pertinent to vision loss rarely observed in other rodent diabetic models (Toh et al. 2019).

The lack of a genome sequence has hindered use of the Nile rat as a model organism. Therefore, we initiated the Nile rat genome project within the framework of the Vertebrate Genomes Project (VGP), which aims to produce reference-quality assemblies of all vertebrate species (Rhie et al. 2021). We present a reference genome of the Nile rat with two complete haplotype-resolved parental genome assemblies. The assemblies are represented by chromosome-scale scaffolds with very few gaps. Over 30 thousand genes and pseudogenes, including sequence-resolved gene duplications, have been annotated. We used this reference genome with additional muroid genomes, in particular the house mouse, for comparative genomics analyses identifying sequences putatively associated with the etiology of diet-induced diabetes. Our findings further demonstrate how haplotype-resolved assemblies and comprehensive gene annotations enable exploration of structural and coding sequence evolution in great detail (C. Yang et al. 2021). This high quality genomic assembly will greatly expand the potential of the Nile rat as a model for biomedical research.

## Results

### The Nile rat assembly is highly complete and highly contiguous

The principal Nile rat individual was sequenced using PacBio continuous long reads for generating contigs, and 10X Genomics linked reads, Bionano optical maps, and Hi-C proximity ligation reads for assembling contigs into scaffolds. Both parents of this individual were sequenced using Illumina short read technology and used to bin the child reads into their respective haplotypes before assembly (Table 1). The assembly, scaffolding, and quality control were performed according to VGP protocols (Rhie et al. 2021). Two sets of haplotype-resolved contigs, paternal and maternal, were generated from PacBio data using TrioCanu (Koren et al. 2018), and scaffolded using 10X Genomics, Bionano Genomics, and Hi-C data (Figure 1b). The paternal haplotype was manually curated to reconstruct and identify chromosomes, and to correct misassemblies and remove false duplications (Howe et al. 2020). The primary pseudohaplotype assembly used for genome annotation consisted of the paternal assembly plus the curated maternal X chromosome. Consistent with the published karyotype (Volobouev et al. 2002), it contained 30 autosomal super-scaffolds and 2 sex chromosomes. In total, the primary assembly contained 2.4 Gb of chromosome-level scaffolds, with an additional 1,534 small unplaced scaffolds. This assembly is highly contiguous (Table 2), with a scaffold N50 of 83 Mbp and a contig N50 of 11 Mbp, one to three orders of magnitude more contiguous than murine genomes assembled using short-reads (Thybert et al. 2018; Lilue et al. 2018) (Figure 1c).

**Table 1.**
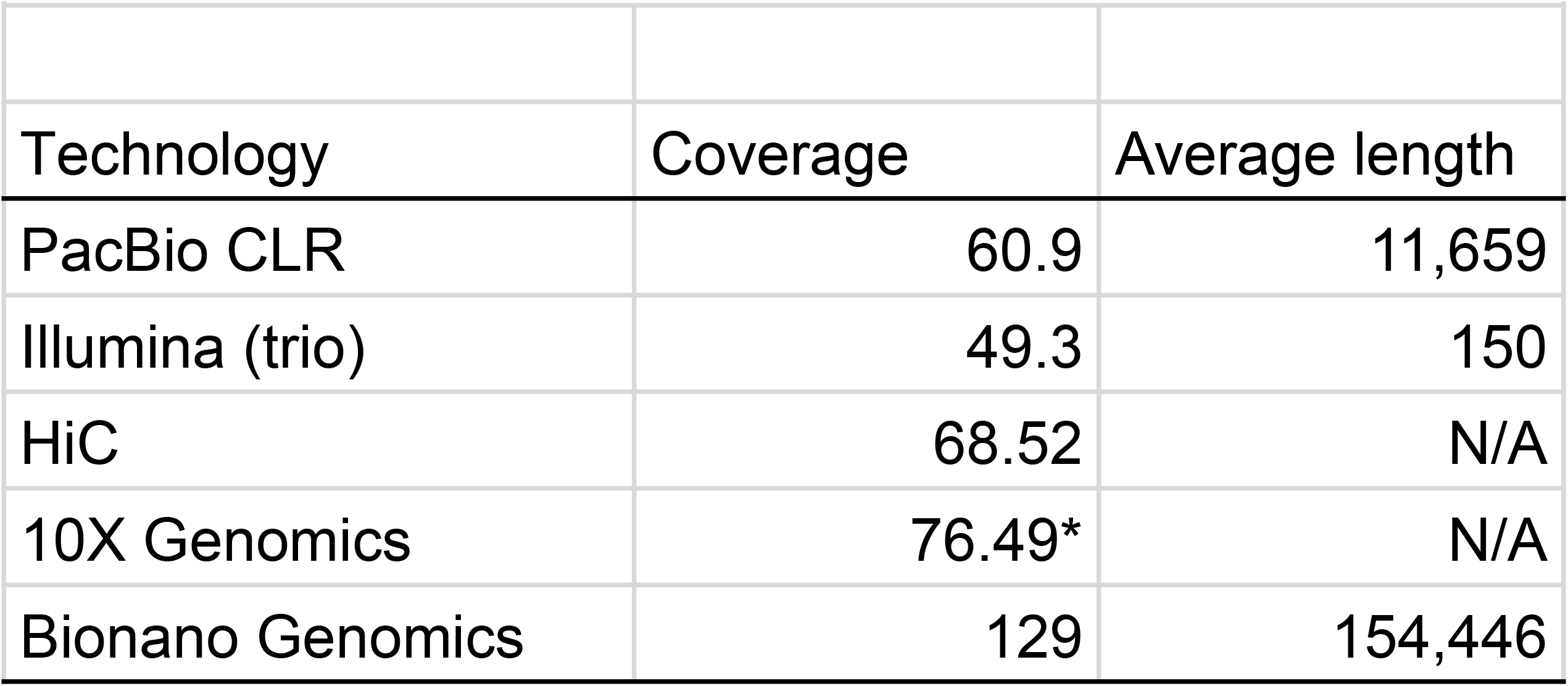
Characteristics of sequencing data. Sequencing coverage and read length of data used to assemble the Nile rat genome. The 10X genomics sequencing coverage is read coverage and not physical (read cloud) coverage. The Bionano genomics coverage counts single-molecule optical maps.

**Table 2.** Assembly statistics.

| **Table 2. Assembly statistics** |  |  |
| --- | --- | --- |
|  | Principal | Maternal |
| Genome size | 2.50 Gb | 2.49 Gb |
| Number of scaffolds | 1,595 | 1,610 |
| Scaffold N50 | 82.7 Mb | 81.2 Mb |
| Number of contigs | 3,219 | 3,135 |
| Contig N50 | 11.1 Mb | 8.9 Mb |
| Percent repeat | 34.4 | 34.32 |
| NCBI protein-coding genes | 22,234 | N/A |
| TOGA gene annotations | 21,038 | 21,284 |

**Figure 1.**
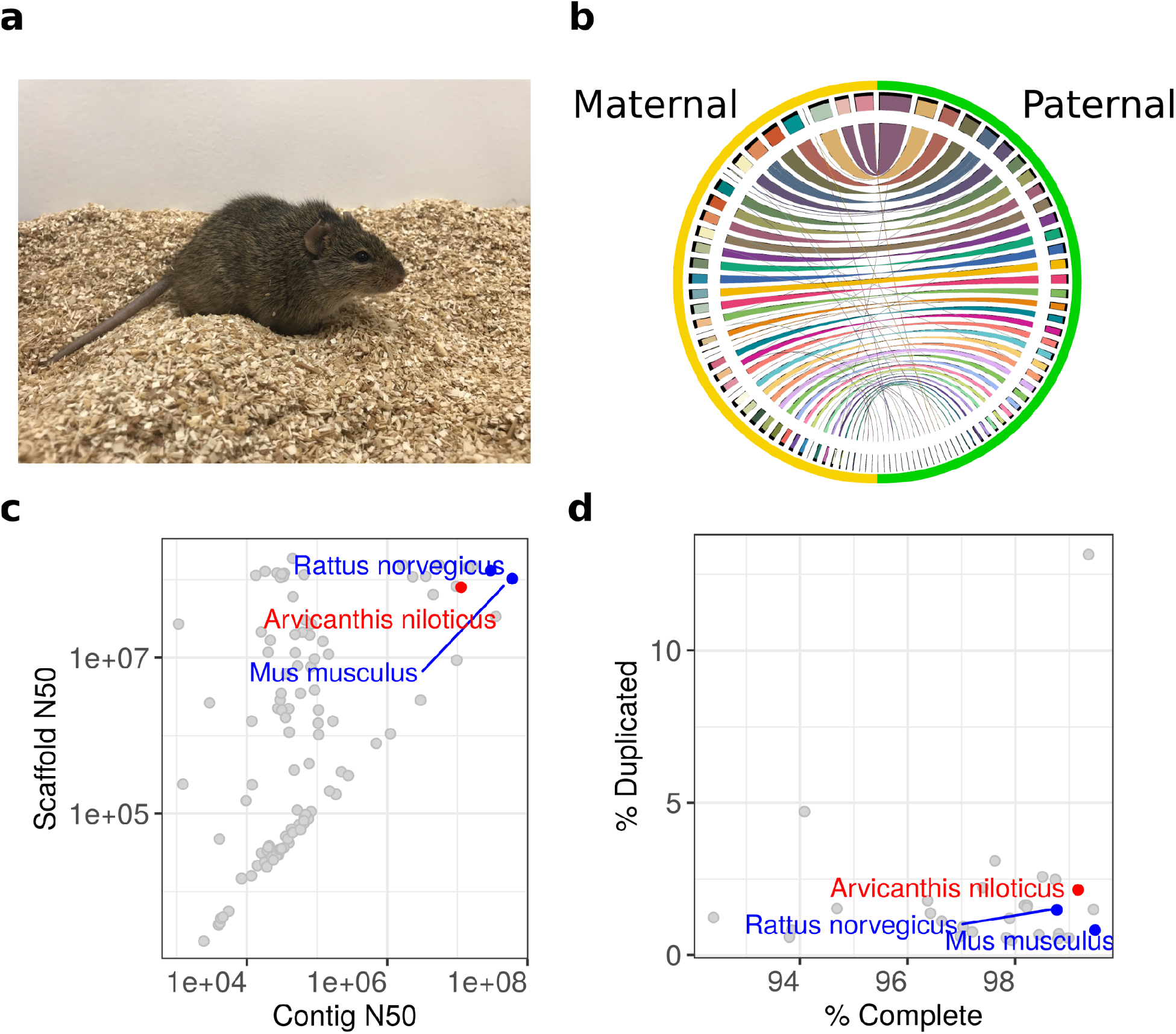
Nile rat genome assembly. **a**. The Nile rat (*Arvicanthis niloticus*). **b**. Scaffolded chromosomes in the maternal and paternal assemblies. Ribbons show similarities between sequences. In order to assess their heterozygosity spectrum, the assemblies have been modified from their GenBank versions as described in Materials and Methods. These modifications are documented in https://osf.io/v4tqc/. **c**. The contig N50 values of Nile rat (*Arvicanthis niloticus*, red), house mouse (*Mus musculus*, blue), Norway rat (*Rattus norvegicus*, blue), and 106 other rodent genomes deposited in GenBank. **d**. Assembly completeness evaluated using BUSCO scores, demonstrating high completeness and average percent duplicated genes that are anticipated to be single-copy genes in rodent genomes.

The BUSCO (Benchmarking Universal Single-Copy Orthologs) annotation references the fraction of genes expected to occur in a single copy in all members of a phylogenetic group, and highlights both the completeness and relative abundance of possible false duplications in an assembly (Kriventseva et al. 2019). The Nile rat BUSCO Complete score is 99% on the Glires subset of OrthoDB version 10 (Kriventseva et al. 2019). We examined the PacBio read depth over duplicated BUSCO genes to see if these duplications were correctly resolved or spurious based on sequencing coverage, setting a permissive threshold of half the average sequencing depth (30.5) to ensure high recall despite fluctuations in mapped coverage. Under this metric, 65% (285/439) are likely correctly assembled. Of the remaining annotated duplications, 63% (97/153) are assembled on unscaffolded contigs and are consistent with higher fragmentation for repetitive sequences known to be problematic for *de novo* assembly. Compared to other rodent genomes, the Nile rat genome assembly has superior contiguity and BUSCO completeness (Figure 1c-d).

A total of 47.2% of the genome is composed of repetitive DNA, as determined using a combination of *de novo* and repeat-library based approaches (Morgulis et al. 2006; Smit, Hubley, and Green 1996), similar to mouse (47.0%) and Norway rat (49.6%) when the same computational pipeline is applied. Repeat content identified by RepeatMasker is summarized in Table 3. The assembly was annotated with the NCBI RefSeq eukaryotic annotation pipeline (Thibaud-Nissen et al. 2013), which identified 31,912 genes and pseudogenes, including 22,234 protein-coding genes. Additionally, we used PhyloPFP (Jain and Kihara 2019) to predict Gene Ontology (GO) terms for all RefSeq proteins (supplementary data: https://osf.io/knmwa/).

**Table 3.** Repeat content of haplotype assemblies. Repeat masking is performed using the rodentia repeat library and RepeatMasker.

| Repeat | Paternal |  | Maternal |  |
| --- | --- | --- | --- | --- |
|  | Number of elements | Percent genome | Number of elements | Percent genome |
| SINE | 1097531 | 6.02 | 1095360 | 6.03 |
| Alu/B1 | 449074 | 1.98 | 447915 | 1.98 |
| B2-B4 | 584908 | 3.81 | 583853 | 3.82 |
| IDs | 34995 | 0.1 | 34992 | 0.1 |
| MIRs | 27964 | 0.13 | 28010 | 0.13 |
| LINEs: | 489920 | 12.26 | 488452 | 12.21 |
| LINE1 | 474324 | 12.15 | 472927 | 12.11 |
| LINE2 | 11535 | 0.08 | 11496 | 0.08 |
| L3/CR1 | 2636 | 0.02 | 2598 | 0.02 |
| LTR | 801475 | 11.23 | 797822 | 11.18 |
| ERVL | 70619 | 0.89 | 70801 | 0.9 |
| ERVL-MaLRs | 355720 | 3.96 | 355437 | 3.99 |
| ERV_classI | 53595 | 0.8 | 52958 | 0.8 |
| ERV_classII | 318728 | 5.55 | 315822 | 5.48 |
| Satellite | 59080 | 0.28 | 58663 | 0.28 |
| Simple repeat | 1289416 | 2.76 | 1284006 | 2.75 |
| Low complexity | 149866 | 0.39 | 149072 | 0.39 |
| Total (including other) |  | 34.4 |  | 34.32 |

### Compilation of type 2 diabetes associated genes

This genome assembly allows us to discover sequence variations that may modify gene functions. Because the Nile rat is an important animal model for type 2 diabetes, we developed a list of genes potentially connected to type 2 diabetes. This list was compiled from gene-disease databases (Davis et al. 2021; Thorn, Klein, and Altman 2013), GWAS catalog from EMBL-EBI (Buniello et al. 2019), and two different text-mining methods (Kuusisto et al. 2020; Raja et al. 2020), resulting in a total of 4,396 genes (Supplementary figure 1). Of these, 3,295 had orthologs identified in the Nile rat assembly annotation by NCBI Orthologs, a part of the NCBI Gene database. The genes of interest were ranked according to the strength of their association with type 2 diabetes. This allowed prioritization of candidate genes in subsequent investigations of genetic variation in the Nile rat genome, including heterozygosity, gene duplication, and positive selection.

### Heterozygosity spectrum of Nile rat, an outbred laboratory species

The Nile rat colony used in this study had been bred in the laboratory since 1998. All individuals were descendants of 29 wild Nile rats from Kenya (McElhinny, Smale, and Holekamp 1997) and have allelic diversity largely reflective of an outbred population. Sequencing a father-mother-offspring trio informs us about genetic heterozygosity (Table 4 and Supplementary Figure 2). Because the paternal and maternal haplotypes were near complete, we were able to investigate heterozygosity more thoroughly than had been possible in most species by comparing homologous chromosomes on the basis of their whole-genome alignment (see Methods). Furthermore, we could compare the numbers of heterozygous variants in Nile rat with those in other mammals. For the principal individual, the rate of single nucleotide variant (SNV) heterozygosity was estimated (by mapping short reads from the same individual) to be 0.086%, about 1/12 of the 1.06% rate estimated across the full spectrum of genetic variants. These estimates are similar to those from the genome assembly of another model organism, the common marmoset, created using a similar VGP trio pipeline, with 0.13% reported for SNV alone and 1.36% for all types of variants (C. Yang et al. 2021). The number of deletions and insertions are approximately equal, as expected, when comparing two haplotypes. We detected 626,683 small deletions (<50 bp) and 4,612 structural variant (SV) deletions (>50 bp), in addition to 626,036 small insertions and 4,632 SV insertions in Nile rat, consistent with other species (Cao et al. 2015; Koren et al. 2018; C. Yang et al. 2021). The distribution of SV by size has several peaks in the length distribution of SVs (Supplementary Figure 3), especially 300 bp, 500 bp in indels and 300 bp, 4.5 kb in other SVs, which matches the common SV sizes of annotated transposable elements, and is consistent with the overall repeat content in the genome (Table 3).

**Table 4.** Genetic variation between the maternal and paternal assemblies.

| Class | Number |
| --- | --- |
| SNV | 2,512,582 |
| Small indel ($\leq 50$ bp) | 1,252,719 |
| Large indel ($> 50$ bp) | 9,244 |
| Inversion | 53 |
| Translocation | 95 |
| Inverted translocation | 62 |
| Copy number variant | 1,971 |

Comparing the two assemblies, we detected 2.51 million SNVs, with 81% of the SNVs confirmed by short-reads. More than one third of all SNVs (862,428) were located within protein-coding genes, and of those, 12,884 SNVs (10,743 SNVs validated by read mapping) were within coding exons. 2,932 SNVs (30%) resulted in nonsynonymous amino acid substitutions affecting 1,581 genes. Iso-Seq data validated 212 of these SNVs in 208 genes, of which 42 were found in our diabetic gene list, including *Alms1* and *Slc19a2*. Human *ALMS1* and *SLC19A2* genes are both involved in monogenic diabetes disorders. Mutations in *ALMS1* can cause Alström syndrome, an autosomal recessive disorder that affects multiple organs where patients typically develop type 2 diabetes in childhood or adolescence (Collin et al. 2002). In the Nile rat, six heterozygous SNVs were validated by testis Iso-Seq data. One missense mutation is *Alms1* 2256P>L, which corresponds to *ALMS1* 3209P>L in humans, scored as probably damaging by PolyPhen and deleterious by SIFT (Flanagan, Patch, and Ellard 2010). Out of 161 mammalian orthologs of *ALMS1*, only three have serine instead of proline in this position, implying that this residue is very well conserved. Similarly, certain *SLC19A2* mutations cause Thiamine Responsive Megaloblastic Anemia syndrome, characterized by diabetes, hearing loss, and anemia (Labay et al. 1999). We found one SNV in Nile rat *Slc19a2*, 275R>W, confirmed by brain Iso-Seq data. This variant is not observed in UniProt, although another variant, 275R>L, is listed in ClinVar as associated with monogenic diabetes with uncertain significance.

### Germline mutation rate

Our trio sequencing data also allows estimating the germline mutation rate. We found four *de novo* candidate mutations in the Nile rat trio, with one mutation of maternal origin and three mutations of paternal origin, suggesting, as in other mammals, a male bias in the contribution to germline mutations. Accordingly, we estimated a *de novo* mutation rate of 0.15 × 10^−8^ mutations per site per generation, though a species estimate requires additional samples.

### Segmental duplications

Long-read assemblies are known to resolve repetitive DNA (Gordon et al. 2016; Rhie et al. 2021). Long, low-copy repeats called segmental duplications (SD) are a class of repetitive DNA that are particularly impactful to phenotypes because they can change gene copy number and reorganize regulatory sequences (Bailey and Eichler 2006). We used a combination of self-alignments (Numanagic et al. 2018) and excess mapped read-depth to quantify SDs in the Nile rat, and compared them against the long-read assemblies of the C57BL strain of house mouse (Sarsani et al. 2019), Norway rat, and white-footed mouse as an outgroup (Long et al. 2019), as well as the mouse reference genome (mm10). Our approach found abundances of SD similar to the existing mm10 annotation, as well as in the long-read assembly of mouse. A total of 123 Mb (4.9% of the genome) of the primary assembly of the Nile rat are annotated as SD, while 114Mb (4.7%) of the maternal assembly are annotated as SD. There are 14.4 Mb of duplications assigned to Y-chromosome scaffolds, indicating there are at least 5.4 Mb of duplicated sequences that differ between parental autosomal chromosomes. Based on excess read depth, 81-106 Mb of additional sequence are missing from the combined diploid assembly due to collapsed duplications.

#### Recent segmental duplication activity in Nile rat

The genomes of all four muroid species have a high proportion of SD with high identity, indicating a recent burst of segmental duplications, consistent with previous observations in the house mouse (Cheung et al. 2003; Thybert et al. 2018) (Figure 2a). For the Nile rat, assuming three generations per year, and *de novo* mutation rate of 0.15 × 10^−8^, roughly 0.5% divergence accumulates every 100,000 years, suggesting that 46% of duplications (64 Mb) are younger than 200k years.

#### Duplicated genes in the Nile rat

We identified gene duplications based on both multi-mapping of entire gene bodies (filtering on sequence identity), and annotation of collapsed copies from excess read depth. To be conservative, multi-mapped isoforms were counted as duplications only if they contained multiple exons spanning at least 1 kb (54k distinct isoforms), with alignments of at least 90% identity and 90% of the original gene length. The high percent identity of most duplicated genes reflects a recent burst of segmental duplications in rodents (Figure 2b). This scheme identified 403 and 369 distinct duplicated genes in the paternal and maternal assemblies respectively; of these, 84/80 were over 99.5% identical, indicating the duplication events occurred within ∼100k years of the present. An additional 13/6 genes are in collapsed regions, indicating there are missing high-identity copies in the assemblies (supplementary data: https://osf.io/4ga9c/), (Vollger et al. 2019).

**Figure 2.**
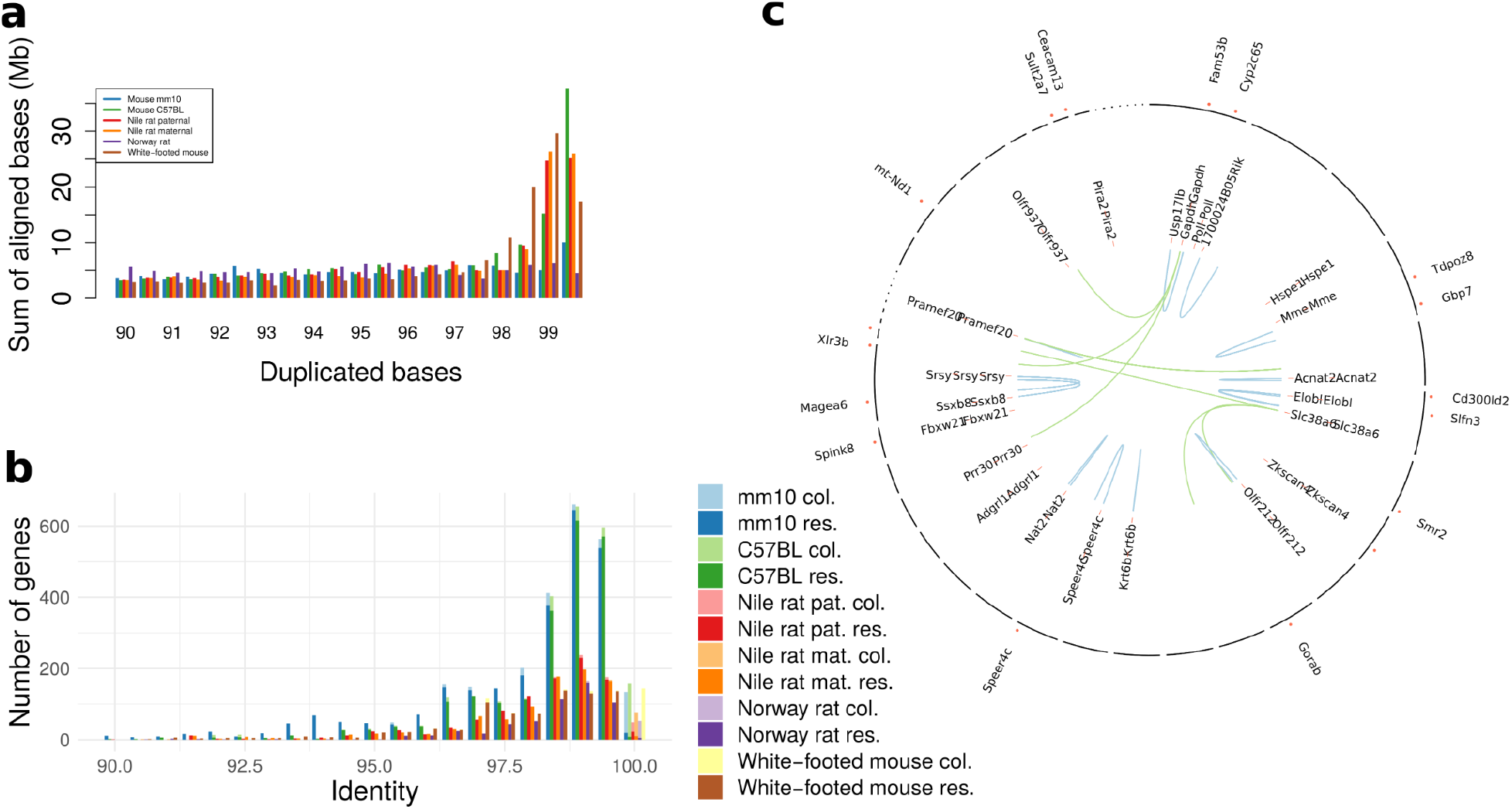
Segmental duplication content in Nile rat and related species. **a**. The total bases annotated as segmental duplication by SEDEF. The total includes all pairwise alignments after filtering for common repeats. **b**. The total number of multi-exon genes duplicated in each of the assemblies for resolved (res.) and collapsed (col.) genes. **c**. Organization of duplicated genes in the Nile rat. Tandemly duplicated genes are in blue; interspersed are in green. Genes in collapsed duplications are indicated as dots in the perimeter. The chromosomes are ordered according to genomic scaffold accessions.

Of duplicated genes with known function, the most common type is olfactory (11.5-18.1%), known to exist as a dense high-copy gene cluster (Sullivan et al. 1996). Additionally, 19.7-20.5% are predicted genes with unknown functions (Supplementary figure 4). Of the remaining duplicated genes in the paternal assembly, 21 are in high-identity duplications (>99.5% identity) with at least 3 copies. Many of these are known to be of high copy number as part of large gene families or mitochondrial genes with many nuclear paralogs. These include the genes counts: *Flg*, 10 copies; *Gapdh*, 6; *Magea*2, 7; *Magea6*, 4; *Pramef* (paralogs *6,17,20*, and *25)*, 4-5; and *Ssxb* (paralogs *1-6,8-10*), 5. The high-quality assembly enables analysis of the mode of expansion of duplicated genes (Figure 2c). For example, some genes were found to have been amplified in tandem arrays, e.g. *Elobl*, 5 copies; *Tdpoz9*, 3; and *Acnat2*, 4 (Figure 3a and b), while others, such as *Slc38a6*, 5 and *Srsy*, 3, are interspersed duplications. The gene *Slfn3* is entirely mapped within a collapsed duplication, with up to three copies missing from the assembly (Figure 3c).

**Figure 3.**
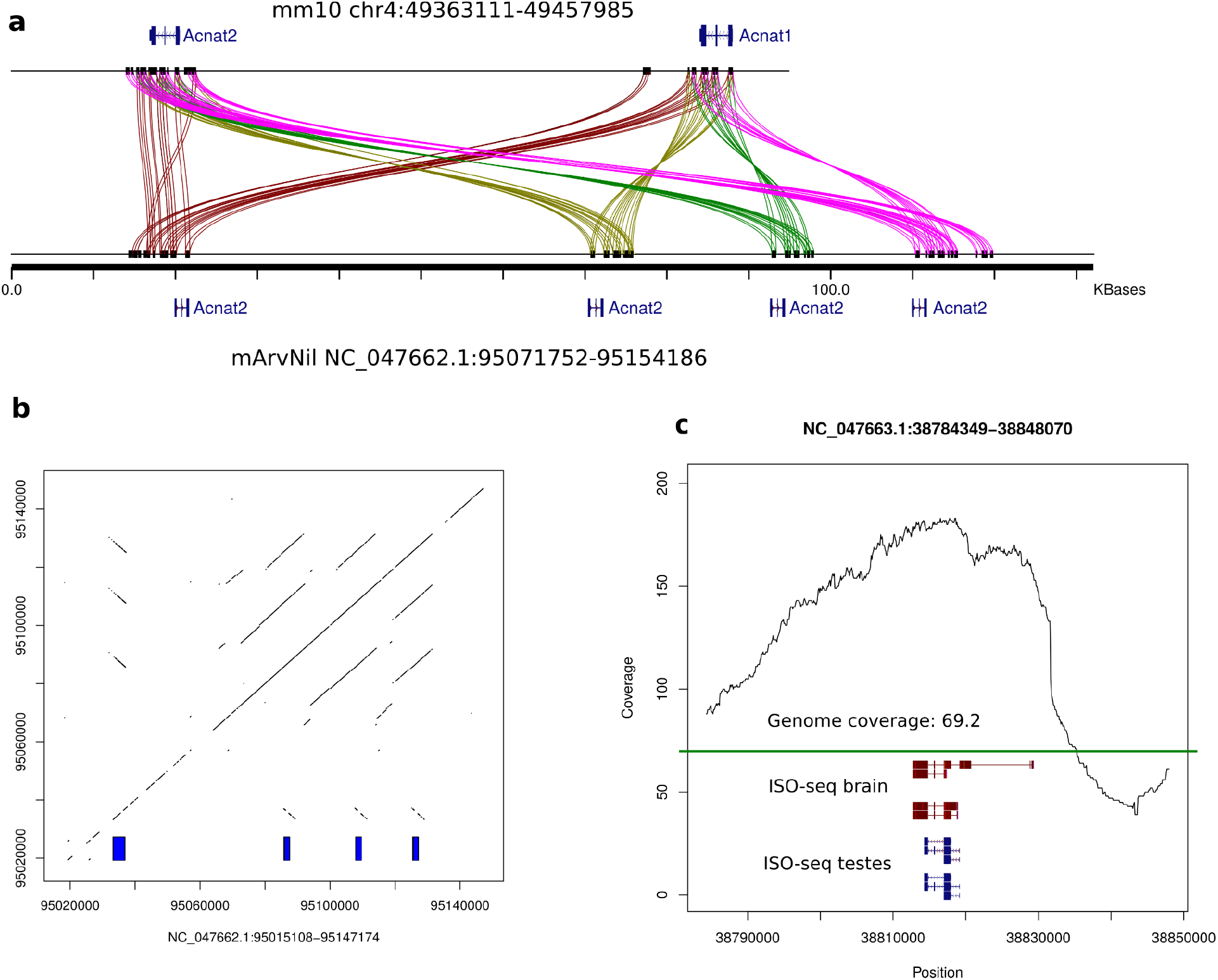
Examples of duplicated genes in Nile rat. **a**. An expansion of the *Acnat2* gene in Nile rat relative to house mouse. Lines are drawn using miropeats (Parsons 1995), with spurious matches outside of gene bodies removed. Colors are used to emphasize gene paralogs. **b**. A dot-plot of the *Acnat2* locus in Nile rat, with gene copies indicated by the blue rectangles. **c**. Read-depth over *Slfn3* in the Nile rat paternal assembly. The average read depth is shown in green, indicating up to four missing copies. The gene is mapped using RefSeq mm10 annotations, with support from PacBio Iso-Seq reads. The gene is associated with immune response, a category of genes that often have large copy-number diversity.

*Acnat* genes arose as duplications of *Baat* genes and likely encode bile acid conjugating enzymes. These genes reside in a highly dynamic locus with multiple gene duplications and gene loss across mammalian species (Kirilenko et al. 2019). Whereas *Acnat* has two copies in the house mouse, our genome reveals four copies in the Nile rat (Figure 3a). This copy number expansion in the Nile rat may affect fatty acid metabolism (Hunt and Alexson 2008) and the synthesis of lipokines (Hernández-Saavedra and Stanford 2019), which may, in turn, have implications for susceptibility to diabetes (Shimoyama et al. 2015).

#### Nile rat has fewer copies of amylase compared to the house mouse

Obesity is a comorbidity with type 2 diabetes (Malik et al. 2010) and has been found to associate with amylase-1 copy number (Falchi et al. 2014). Individuals from human populations with high-starch diets tend to have more copies of *Amy1* (Perry et al. 2007). The amylase locus in the house mouse genome contains seven protein coding genes -*Amy1*, five copies of *Amy2a* (*Amy2a1-Aym2a5*), and *Amy2b*, as well as one amylase-like pseudogene. Our approach to annotating resolved gene duplications did not detect duplicated copies of amylase in Nile rat.

However, RefSeq annotations, as well as TOGA projections of human and mouse genes to the Nile rat genome, annotate a cluster of three protein coding amylase genes, two amylase-2 andone amylase-1, plus two amylase-like pseudogenes (Figure 4a). The two amylase-2 genes share 81.7% sequence identity across the genomic intervals, while the mouse amylase copies range from 98.6 to 99.9 percent identity with each other (Figure 4b). Furthermore, a multiple sequence alignment of amylase proteins from Nile rat and three other rodents, house mouse, Norway rat, and white-footed mouse, shows the Nile rat amylase-1 clustering with amylase-1 proteins of the other three species, while amylase-2 copies from each species form separate clusters (Figure 4c). This indicates that multiple amylase-2 genes are the result of recent expansions that happened independently in different muroid lineages. The two amylase-2 copies in Nile rat are more divergent from each other than any two of the four full length copies in the house mouse, perhaps reflecting the latter’s recent adaptation to commensalism with humans.

**Figure 4.**
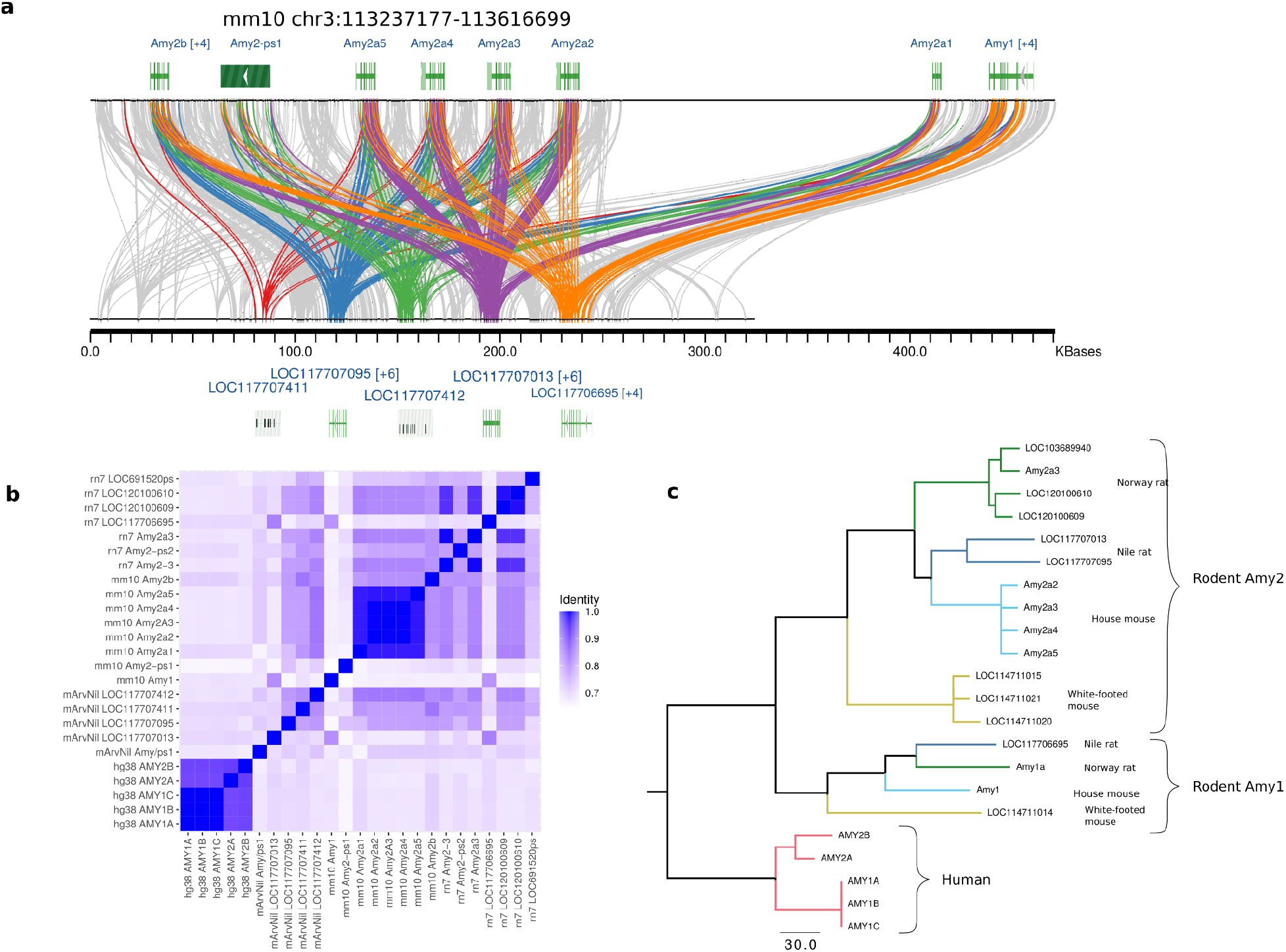
**a**. The sequence homology in the amylase locus for mouse (top) and Nile rat rendered using miropeats. The five TOGA annotations of amylase are each rendered using separate colors. The blue and purple copies show amylase-2 homologies, orange is amylase-1, and red/green lines are annotated pseudogenes. **b**. Pairwise similarity of amylase genes in human, Nile rat, mouse, and Norway rat, ordered according to their genomic coordinates. **c**. A phylogenetic tree using COBALT multiple sequence alignment of amylase proteins from each of the four genomes in (B) and white footed mouse.

#### Extensive duplication of Ybx3-like retrogenes

Y-box binding proteins are a major group of cold shock proteins defined by the presence of a cold shock domain (CSD), which has DNA-and RNA-binding capabilities. Mammals have a family of three paralogous Y-box binding proteins - *Ybx1, Ybx2*, and *Ybx3*. Among its diverse biological roles, *Ybx3* is involved in nutrient sensing, a function commonly dysregulated in metabolic diseases. It controls the intracellular levels of large neutral and aromatic amino acids (Cooke et al. 2019), including the branch chained amino acids highly associated with insulin resistance and obesity (White et al. 2021). The NCBI genome annotation pipeline found 56 “Y-box-binding protein 3-like” (*Ybx3-like*) genes and pseudogenes in the Nile rat genome, 26 of which were annotated as protein-coding, while a BLAT search of the canonical *Ybx3* transcript against the genome found 147 hits, all dispersed throughout the genome (Figure 5a).

**Figure 5.**
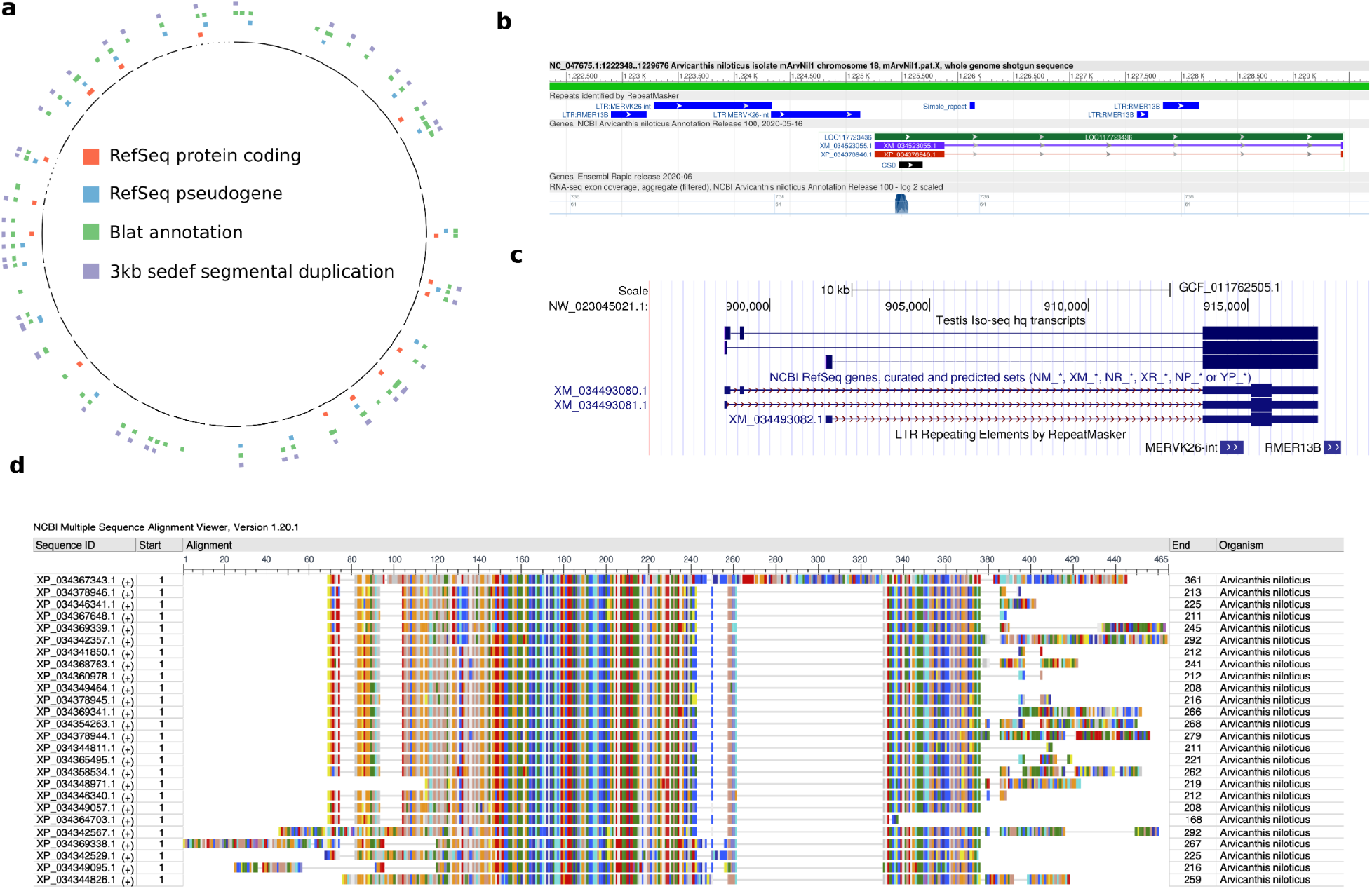
Ybx3-like elements in the Nile rat genome. **a**. Ybx3-like elements are interspersed throughout the genome. Many have been annotated as protein coding genes or pseudogenes by NCBI. Most are recognized as segmental duplications by SEDEF. **b**. The architecture of a typical Ybx3-like gene, *LOC117723436*, visualized in the NCBI Genome Data Viewer. This gene has one large and one small exon. The large exon is flanked by MERVK26-int and RMER13B endogenous retroviral elements. It contains a CSD domain and is partially supported by short read RNA-seq data. **c**. Expression of *LOC117701283* in testis visualized in the UCSC genome browser. The three Iso-seq transcripts have identical CDSs, represented by thick boxes. MERV26-int is located in the 5’ UTR region, rather than outside the large exon like in most other *Ybx3-like* genes. **d**. Multiple alignment of predicted *Ybx3-like* proteins and the canonical *Ybx3*, visualized by NCBI COBALT. This visualization uses the Rasmol color scheme, where amino acids with similar properties are shown in matching colors. The canonical protein is in the first row.

*Ybx3-like* genes consist of a single large exon and often one or two small exons. The large exon is consistently flanked by two endogenous retroviral elements (ERVs), MERVK26-int upstream and RMER13B downstream, often with more than one copy of each (Figure 5b). SEDEF annotated 78 segmental duplications that map to these genes, averaging 3.3 kb in length. These duplications encompass the large exon and the flanking ERVs. The large exon of all *Ybx3-like* genes annotated as protein-coding by RefSeq contains the CSD. One *Ybx3-like* gene, LOC117701283, is supported by three full length transcripts found in our testis Iso-seq dataset (Figure 5c). The canonical *Ybx3* in Nile rat has 9 exons. An alignment of the predicted *Ybx3-like* proteins to the protein product of the canonical gene shows that their large exon contains most of the canonical sequence, with the exception of a large gap and the C-terminal region (Figure 5d). The gap contains a portion of exon 5 and most of exon 6 of the canonical protein and the missing N-terminal region contains a portion of exon 8 and the entire exon 9. Many *Ybx3-like* proteins also contain N-terminal and/or C-terminal segments that are not homologous to canonical *Ybx3*.

#### Duplicated Nile rat genes that exist as single copy genes in house mouse

Gene copy numbers often vary both between species and between individuals within a species. We identified 117 genes that had two or more copies in both Nile rat haplotypes while being present in a single copy in both house mouse assemblies. Of the genes duplicated in Nile rat but not in house mouse, 21 were in our diabetic gene set including *Gckr* and *Fndc4. Gckr* encodes glucokinase regulatory protein, which forms an inhibitory complex with glucokinase thereby regulating uptake and storage of dietary glucose (Raimondo, Rees, and Gloyn 2015). Mammalian *Gckr* is composed of two sugar isomerase (SIS) domains which contain binding sites for fructose-6-phosphate (F6P) or fructose-1-phosphate (F1P) and glucokinase, where the fructose metabolites alter the affinity of *Gckr* for glucokinase (Sanghera et al. 2019). Human *GCKR* has been reported as a diabetic susceptibility gene by several studies (Vaxillaire et al. 2008) (Sparsø et al. 2008) (Diabetes Genetics Initiative of Broad Institute of Harvard and MIT, Lund University, and Novartis Institutes of BioMedical Research et al. 2007) (Zahedi et al. 2021) (Chen et al. 2021). We found a full-length second copy of *Gckr*, LOC117716845, 1.2 Mb downstream of the canonical *Gckr*. This copy of *Gckr* has been annotated as a protein coding gene by Ensembl, ENSANLG00005017071, but RefSeq annotated it as a pseudogene. It is a high identity copy (96.7%) and both SIS domains are present. However, the F1P binding site in the first SIS domain (Pautsch et al. 2013) is affected by non-synonymous substitutions (Supplementary Figure 5). Additionally, a BLAT search of the Nile rat canonical *Gckr* transcript yielded 25 hits. Among them, there were four other, truncated copies of high identity (92-94%) multi-exon glucokinase regulator protein-like pseudogenes. We did not detect any *Gckr* duplications in white-footed mouse or Norway rat, suggesting that this is a copy number gain in Nile rat, rather than a loss in the house mouse. A similarly duplicated gene found in Nile rat but not house mouse is *Fndc4*, where the second copy is almost full-length and validated by testis Iso-Seq data, with 97.8% identity and is located 1.3 Mb downstream of the canonical *Fndc4*. Fndc4 attenuates hyperlipidemia-induced insulin resistance in mice (Lee et al. 2018).

### Differences in protein coding gene content between Nile rat and house mouse

We used TOGA (Kirilenko et al., submitted) to project protein coding genes from human and house mouse to the Nile rat genome. Overall, 99.7% of TOGA annotated genes in the paternal assembly are also annotated in the maternal assembly using mouse gene models; when using human gene models the number is 96.6%.

We compared TOGA projections from the mouse to genes predicted by the NCBI genome annotation pipeline and explored the genes that differ between the two species. 516 mouse genes appeared to be missing from Nile rat, in so far as TOGA was not able to project them to the primary haplotype assembly (supplementary data: https://osf.io/gbtws/). However, the majority of these genes were present in the assembly of the other haplotype. Examination of two examples that were part of our diabetic gene list revealed that structural variation and assembly errors were the primary reasons to not identify a gene. For example, *Hadh* is partially disrupted due to a gap in the primary haplotype assembly of the Nile rat but appears intact in the alternate (Supplementary Figure 6). A different case is represented by orosomucoid 2, *Orm2*. Orosomucoid is a potential diabetes biomarker (Tsuboi et al. 2018). A cluster of four *Orm* genes in house mouse, including *Orm1, Orm2, Orm3*, and the pseudogene *Gm11212*, corresponds to a single gene in Nile rat, annotated by RefSeq as *Orm1*. TOGA has mapped *Orm1* and *Orm3*, but not *Orm2*, to the primary haplotype of the Nile rat. Cactus alignments confirmed the existence of a four-fold duplication in house mouse compared to Nile rat in this locus in both Nile rat haplotypes (Supplementary Figure 7). Conversely, 1,601 Nile rat genes annotated by NCBI could not be projected to mouse genes (supplementary data: https://osf.io/92eu7/). Most were members of families of duplicated genes, including retrogenes derived from *Ybx3* and ribosomal proteins. A gene set enrichment analysis of these genes is discussed in the Supplement (Supplementary Figure 8).

There were 218 mouse genes that TOGA was unable to map to either of the Nile rat haplotypes (supplementary data: https://osf.io/vd5jn/), ten of which were in our diabetes gene list (supplementary data: https://osf.io/ahvmn/). The two top-ranked of these, *Hmga1b* and *G6pd2*, are retrogenes that have emerged in the mouse lineage from parental genes *Hmga1* (Foti et al. 2005) and *G6pd (Heymann, Cohen, and Chodick 2012)*, respectively (Supplementary Figures 9 and 10).

Conversely, 69 genes were absent in the house mouse and present in both Nile rat haplotypes. Seven of these genes were in our diabetic gene list (supplementary data: https://osf.io/84xvp/) including *Aqp10*. Although *Aqp10* is a protein coding gene in the Nile rat and human, it is present as a nonfunctional pseudogene in the mouse (Morinaga et al. 2002). Human *AQP10* has been suggested to be a target for obesity and metabolic diseases (Gotfryd et al. 2018), but could not be studied in the house mouse where the gene has been pseudogenized.

### Positively selected genes

We identified 119 positively selected protein coding genes in Nile rat, comparing it with eight other species in the *Myomorpha* suborder via the branch-site model implemented in PAML (v4.9j), using human as an outgroup (Supplementary Table 1, Supplementary Data https://osf.io/3xqh6/). To avoid confounding effects dependent on assembly quality and isoform differences, protein coding genes from all species were re-annotated using exonerate v2.4 (Slater and Birney 2005). After filtering, 7,492 high quality orthologous genes remained in this dataset (Gudmundsson et al. 2021). Out of these genes, 26 had human orthologs previously found to have low tolerance to mutation, where ≤ 20% of expected loss-of-function variants were observed in population-scale exome sequencing data annotated by gnomAD (Gudmundsson et al. 2021). Of these 26 genes, *Xiap, Ppp2r5e, Krt1, Pik3r5*, and *Irf5* had amino acid substitutions in the Nile rat that did not exist in any other rodents with NCBI annotated genomes.

Here, we take a closer look at two of these genes. X-linked inhibitor of apoptosis protein (*XIAP*) prevents apoptosis of islet β-cells and is considered as a therapeutic target against β-cell destruction in diabetes (Plesner et al. 2005). *XIAP* is strongly intolerant to sequence variations, with only 2 out of 16.3 expected loss-of-function SNVs observed in humans (Gudmundsson et al. 2021). In the Nile rat, we found three sites that were under positive selection, at positions 122, 135, and 190 within the protein sequence. Residues 135 and 190 are well-conserved across mammalian genomes. The absence of human variants in corresponding positions and the presence of nearby disease variants could indicate that mutations of these residues are consequential (Supplement). Like *XIAP*, Protein Phosphatase 2 Regulatory Subunit B’Epsilon, *PPP2RE*, is also a diabetic gene associated with pancreatic islets (Pedersen, Gudmundsdottir, and Brunak 2017). In humans, loss-of-function variants of *PPP2R5E* have not been reported, and only 31% of expected missense SNVs were observed (Gudmundsson et al. 2021). In the Nile rat, we found a *Ppp2r5e* 269 I>L substitution, whereas isoleucine at this position is universally conserved across all other mammals (supplementary data https://osf.io/sdyuh/).

## Discussion

The Nile rat is an important animal model for studying type 2 diabetes and for biomedical research dependent on a diurnal chronotype. We have generated a highly contiguous, haplotype-resolved genome assembly of this species. A haplotype-resolved assembly can enable a more complete annotation by virtue of having two distinct assemblies to work with. For example, an incomplete *Hadh* gene in the paternal assembly is resolved in the maternal assembly. While the BUSCO duplication value appears to be higher than many other rodent assemblies, a read-depth analysis suggests that 65% of redundant BUSCOs are likely actual duplicated genes, indicating that the BUSCO gene annotations may be improved as additional high-quality genomes are analyzed.

A haplotype-resolved assembly enabled us to explore all types of heterozygosity, including SNVs and SVs (indels, and other structural polymorphisms of all sizes). Overall, we observed a level of heterozygosity consistent with an outbred organism. However, chromosomes 1, 3, and 5 had large regions of low heterozygosity. These may have resulted from inbreeding of close relatives that occurred at generation 4 due to small colony size, although no direct brother-sister matings were used. The sequenced individual is from generation 6.

Because a trio pedigree was sequenced for the Nile rat, we were also able to calculate the rate of *de novo* germline mutations. This rate is 0.15 × 10^−8^ mutations per site per generation, lower than the mutation rates reported for other mammals (Bergeron et al. 2021). A more accurate, population-based estimate can be a subject of future research.

Our high-quality assembly enabled us to resolve most segmental duplications and catalog multicopy genes. However, some collapses remain, which encompass 6-16% of all multicopy genes. While advances in technology and assembly algorithms will reduce the number of collapsed duplications, annotations of genes in collapses should be continued until an approach guaranteeing telomere-to-telomere assembly is established.

A comparative analysis of the Nile rat with other rodents enabled us to detect several types of evolutionary events affecting genes known to be involved in the mechanisms of type 2 diabetes (Table 5). We selected ten genes for a closer examination. *Gckr* (Raimondo, Rees, and Gloyn 2015), *Fndc4 (Frühbeck et al. 2020), amylase*, and *Orm* are differently affected by segmental duplications in Nile rat and house mouse. *Hmga1b* and *G6pd2* have been created by retrotransposition in the mouse. Multiple copies of *Ybx3-like* genes have been created in Nile rat by retrotransposition followed by segmental duplication, similar to *TP53* in elephants (Sulak et al. 2016). *Alms1* and *Slc19a2* are affected by heterozygosity, and *Xiap* by positive selection in the Nile rat lineage.

**Table 5.** Copy number divergent, heterozygous, and positively selected genes.

|  |  |  |  |
| --- | --- | --- | --- |
| Category | Number of genes | Linked to type 2 diabetes | Comment |
| Human-house mouse orthologs | 18,951 | 3,567 | Based on the MGI orthologs table |
| Human-Nile rat orthologs | 16,235 | 3,295 | Based on the NCBI Gene database |
| Duplicated in mouse but not in Nile rat | 368 | 15 |  |
| Duplicated in Nile rat but not in mouse | 117 | 21 |  |
| Mouse genes missing in Nile rat | 218 | 10 | Homozygous-missing only |
| Named Nile rat genes missing in mouse* | 69 | 7 | See footnote |
| Non-synonymous SNVs supported by Iso-seq | 208 | 42 |  |
| Positively selected | 119 | 19 |  |
| *Assigned names other than locus numbers by NCBI, no mouse genes mapped by TOGA, and names do not occur in the MGI table of human-mouse orthologs |  |  |  |

Retrotransposition and segmental duplication are major drivers of genome evolution, including creation of new genes. Comparing reference-quality assemblies of closely related species enabled us to observe these events at high levels of detail. The ability of these new genes to express functional proteins and their impacts on the biology of the Nile rat must be the subjects of future studies. We hope that the availability of a reference-quality genome of this important species will both inspire and enable future research.

## Materials and Methods

### Nile rat tissue collection

Spleen, brain, and testis tissue were collected from a 21-week-old male Nile rat (T564M) in the laboratory colony of Huishi Toh and James Thomson at University of California, Santa Barbara. The spleen was used for genome sequencing, whereas the brain and testis were used for transcriptome analysis. These tissue samples were flash frozen in liquid nitrogen immediately after dissection. Additionally, blood samples were collected via cardiac puncture from Nile rat T564M’s parents (T480F and T469M). T564M is a generation 6 descendent of 17 breeders that were imported from the Hayes lab in Brandeis University, a secondary colony of the original laboratory colony from the Smale lab in Michigan State University, which started from 29 wild Nile rats captured in Kenya. The Nile rats in this study were fed a high fiber diet 5326 and were normoglycemic. All animal experiments were approved by the University of California, Santa Barbara, Institutional Animal Care and Use Committee, and conducted in accordance with the NIH Guide for the Care and Use of Laboratory Animals.

### Genome sequencing

#### Primary subject

We isolated 23 ug of ultra high molecular weight DNA (uHMW) from 35 mg of flash-frozen spleen tissue using the agarose plug Bionano Genomics protocol for animal tissue (DNA isolation fibrous tissue protocol #30071C). uHMW DNA quality was assessed by a Pulsed Field Gel assay and quantified with a Qubit 2 Fluorometer. 10 μg of uHMW DNA was sheared using a 26 G blunt end needle (Pacbio protocol PN 101-181-000 Version 05). A large-insert Pacbio library (CLR) was prepared using the Pacific Biosciences Express Template Prep Kit v1.0 (#101-357-000) following the manufacturer protocol. The library was then size selected (>20 kb) using the Sage Science BluePippin Size-Selection System. The Pacbio library was sequenced on 22 PacBio 1M v3 (#101-531-000) SMRT Cells on a Pacbio Sequel instrument using the sequencing kit 3.0 (#101-597-800) and a 10 hour movie. A total of 206.97 Gb of raw reads data with an average insert size N50 of 23,715 bp bases was generated. Unfragmented uHMW DNA was used to generate a linked-reads library on the 10X Genomics Chromium (Genome Library Kit & Gel Bead Kit v2 PN-120258, Genome Chip Kit v2 PN-120257, i7 Multiplex Kit PN-120262).

From this 10X library, we generated 256.78 Gb of sequence data on an Illumina Novaseq S4 150bp PE lane. uHMW DNA was labeled for Bionano Genomics optical mapping using the Bionano Prep Direct Label and Stain (DLS) Protocol (30206E) and run on one Saphyr instrument chip flowcell. Hi-C preparation was performed by Arima Genomics using the Arima-HiC kit (P/N: A510008), and an Illumina-compatible library was generated using the KAPA Hyper Prep kit (P/N: KK8504). This library was then sequenced on an Illumina HiSeq X (150bp PE) at 129X coverage following the manufacturer’s protocols. Sequencing read lengths and depths of coverage are summarized in Table 1.

#### Parents

PCR-free Illumina libraries were generated from 1 μg genomic DNA using a Covaris LE220-plus to shear the DNA and the TruSeq® DNA PCR-Free HT Sample Preparation Kit (Illumina) for library generation. The median insert sizes were approximately 400 bp. Libraries were tagged with unique dual index DNA barcodes to allow pooling of libraries and minimize the impact of barcode hopping. Libraries were pooled for sequencing on the NovaSeq 6000 (Illumina) to obtain at least 750 million 151-base read pairs per library. This resulted in 49.3X coverage of the parental genomes.

### Transcriptome sequencing

We extracted and purified total RNA from brain and testis tissues using the QIAGEN RNAeasy kit (Cat. No. 74104). For each tissue, 25-30 mg was cut into 2mm pieces before homogenization with the Qiagen TissueRuptor II (Cat No./ID: 9002755). The quality of all RNAs were assessed using a Fragment Analyzer (Agilent Technologies, Santa Clara, CA) and quantified with a Qubit 2 Fluorometer (Qubit™ RNA BR Assay Kit - Catalog number: Q10210).

PacBio Iso-seq libraries were prepared according to the ‘Procedure & Checklist - Iso-Seq Template Preparation for Sequel Systems’ (PN 101-070-200 version 05). Specifically, cDNA was reverse transcribed using the SMRTer PCR cDNA synthesis kit (Clontech, Mountain View, CA) from 329 ng and 374 ng of total RNA for brain and testis respectively. Amplified cDNA was cleaned with AMPure beads and a PacBio library was prepared using the Pacific Biosciences Express Template Prep Kit v1.0 (#101-357-000) following the manufacturer protocol. PacBio Iso-seq libraries were sequenced on a PacBio Sequel (sequencing chemistry 3.0) with 20 hours of movie time. We sequenced one SMRT Cell for each Iso-seq library. We then used the Iso-seq application in the PacBio SMRT Link package to generate Circular Consensus Sequences (CCSs), remove cDNA primers and concatemers, identified strandedness, trim polyA tails, and perform *de novo* clustering and consensus call to output high-quality full-length consensus isoforms.

### Genome assembly and annotation

The haplotype-resolved assembly was generated using TrioCanu v. 1.8 using the parental Illumina reads and the PacBio WGS data (Koren et al. 2018). Consensus sequences were generated using Arrow v. smrtlink_6.0.0.47841 (Pacific Biosciences), followed by purging of spurious duplications using purge_dups v. 1.0.0 (Guan et al. 2020). The assemblies were then scaffolded using 10X Genomics linked long reads with scaff10x v. 4.1.0, Bionano optical maps with Solve v. 3.2.1_04122018, and HiC data with Salsa2 HiC v. 2.2. The scaffolds were polished using PacBio reads with Arrow and 10X Genomics synthetic long reads with Longranger and Freebayes v. 1.3.1. This was followed by decontamination and manual curation (Howe et al. 2020). The mitochondrial genome was assembled using mitoVGP workflow v2.0 (Formenti et al. 2021).

The genome was annotated using the RefSeq eukaryotic annotation pipeline (O’Leary et al. 2016) with 73,241 brain and testes Iso-Seq full-length transcript sequences (Wang et al. 2016). There were 457,991 isoforms in 21,723 distinct coding regions. The quality of the consensus was sufficiently high that the majority of annotated gene models were complete; 2.7% of genes (591/21,723) required modification of the reference to account for frameshift errors.

We used Phylo-PFP (Jain and Kihara 2019) to assign Gene Ontology (GO) terms to protein coding genes. Phylo-PFP is a sequence-based protein function prediction method which mines functional information from a broad range of similar sequences, including those with a low sequence similarity identified by a PSI-BLAST search. The sequences retrieved from PSI-BLAST are reranked by considering the phylogenetic distance and the sequence similarity to the query. Incorporating phylogenetic information leads to better functional similarity estimation. Gene Ontology (GO) terms of each retrieved protein are assigned the same score as the sequence. Finally, for each GO term, scores from all sequences are summed. The prediction is also enriched with GO terms that have greater than 90% probability of co-occurrence.

### Assembly quality metrics

In order to evaluate the quality of our assembly, we compared it to representative genomes of other species of rodents available from the NCBI assembly database. We utilized R package rentrez, a wrapper for NCBI E-utilities, to retrieve assembly records. The R script used for the retrieval and plotting of assembly quality metrics is available on OSF at https://osf.io/r9uqf/.

### Segmental duplication analysis

The annotation pipeline is available at https://github.com/ChaissonLab/SegDupAnnotation/releases/tag/vNR

#### Segmental duplication annotation with self alignments

Genomes were repeat masked using the union of windowmasker v1.0.0 and RepeatMasker 4.1.1 with the parameter “-species rodentia”. An initial set of segmental duplications were identified using SEDEF version 1.1-37-gd14abac-dirty with default parameters (Numanagic et al. 2018), that were then filtered in post-processing to remove mobile elements annotated as segmental duplications. First, duplications were removed if either copy was over 90% repeat masked. Next, the remaining annotations contained duplications that were 1-2 kb, high copy (>20 copies), and were typically masked as endogenous retroviruses using the CENSOR repeat masking server (https://www.girinst.org/cgi-bin/censor/censor.cgi). To remove these, high-copy duplications were detected and filtered from the duplication set. The multiplicity of a duplication was measured considering transitive copies potentially missed in alignments by creating a graph where every repeated interval corresponds to a node, and edges connect both the pair of nodes corresponding to the repeat alignments, and any overlapping intervals. The number of unique intervals in each connected component was used to assign a repeat copy number, and repeats with copy number greater than 20 were removed.

#### Gene duplication annotation

Duplicated genes were annotated using multi-mapped sequences. Gene models were defined using Nile rat RefSeq sequences aligned using minimap2 using the −x splice option. Next, sequences of genes with at least one intron with a gene body of at least 1 kb were mapped back to each assembly using minimap2. Alignments with at most 10% divergence that were at least 90% of the query sequence length were considered as duplicated genes. A single isoform for each gene was retained as a duplication. When multiple genes map to the same location, only the first sequence mapped by the pipeline is retained. The number of copies of a gene are counted in the resulting set of alignments.

#### Annotation of collapsed repeats

We used a hidden Markov model to assign copy numbers to collapsed duplications. Each copy number is encoded as a hidden state from 0 to a maximum of 12 copies. The observed data are the coverage values in 100-base bins across each assembly. The probability of emission is calculated as a negative binomial with a mean and variance estimated according to the copy number of each state based off of the mean observed at the copy-number two sites in the genome.

### Mutation rate analysis

The offspring and parental reads were mapped to each assembly independently (paternal and maternal). Duplicate reads and reads mapping to more than one region were removed. Variants were called using GATK 4.0.7 HaplotypeCaller in base-pair resolution mode, calling each single site of the genome. Two independent joint genotyping analyses were carried out: one for the three individuals (mother, father, offspring) mapped to the maternal assembly and one for the three individuals mapped to the paternal assembly. The variant file was filtered on the quality of the genotyping features following these parameters: QD < 2.0, FS > 20.0, MQ < 40.0, MQRankSum < −2.0, MQRankSum > 4.0, ReadPosRankSum < −3.0, ReadPosRankSum > 3.0, SOR > 3.0.

Additional filters were applied at each position to detect the candidate mutations. Thus, a site would be filtered out if one individual had:

- a depth DP < 0.5 × depth _individual_ and DP > 2 × depth _individual_, with depth _individual_ being the average depth of the individual (depth _individual_ offspring: 56 X, depth _individual_ father: 78 X and depth _individual_ mother: 84 X).
- a genotype quality GQ < 60.
- a number of alternative alleles in the parents with AD > 0.
- an allelic balance in the offspring with AB < 0.3 and AB > 0.7.

We then identified the maternal *de novo* candidates using the following genotypes:

- sites where the parents are homozygous for the reference (0/0) and the offspring is heterozygous (0/1) when mapped to the paternal genome: 35 candidates
- sites where the parents are homozygous for the alternative (1/1) and the offspring is heterozygous (0/1) when mapped to the maternal genome: 133 candidates

A comparison of the reads in the candidates sites resulted in only one position with an overlap of read names. Thus, we found one maternal *de novo* candidate mutation.

Similarly, we identified the paternal *de novo* candidates using the following genotypes:

- sites where the parents are homozygous for the reference (0/0) and the offspring is heterozygous (0/1) when mapped to the maternal genome: 38 candidates
- sites where the parents are homozygous for the alternative (1/1) and the offspring is heterozygous (0/1) when mapped to the paternal genome: 143 candidates

The comparison of reads in both datasets resulted in three positions with overlapping reads. Thus, we retained three paternal *de novo* candidate mutations.

To estimate a per generation rate, we calculated callability, the number of sites with full detection power. These were all the sites that passed the DP, the GQ, and the AD filters. The maternal callability was 1,371,536,436 base pairs and the paternal callability was 1,365,805,112 base pairs. This callability estimation does not take into account the filters applied only on polymorphic sites that could have reduced the detection power on some of the callable sites. To correct for any bias due to the site filters and the allelic balance filter we applied a false negative rate (FNR) correction on the callability. The FNR was calculated as the number of true heterozygous sites, i.e. one parent homozygous for the reference allele, one parent homozygous for the alternative allele and the offspring heterozygous, filtered out by the AB filter. This FNR also took into account the proportion of callable sites expected to be filtered out by the site filters if a variant was present. FNR was ∼5% on both the maternal and paternal assembly. Finally, we estimated the mutation rate using a diploid genome size of 2.6 Gb.

### Heterozygosity spectrum

To call heterozygous sites between the two haploid sequences, we directly compared two haploid assemblies using Mummer (v3.23) with the parameters of “nucmer -maxmatch -l 100 -c 500”. Before retrieving all spectrum of genetic variants, we refined haplotype genomes by anchoring the scaffolds which might be lost in final assemblies (Supplementary Figure 11). SNV and small indels were generated by “delta-filter -m -i 90 -l 100” and followed by “dnadiff”. Several custom scripts were used to deal with Mummer output (supplementary data: https://github.com/comery/Nile_rat). We employed Assemblytics v1.2 (Nattestad and Schatz 2016) and SyRi v1.0 (Goel et al. 2019) to detect SVs from Mummer alignment using default parameters. Specifically, Assemblytics for large indels and CNV and SyRi for inversions, translocations, and other SVs. SVs in which more than half the feature consisted of gaps were dropped.

### Branch-site test analysis

To find positively selected genes (PSGs) in the Nile rat lineage, we compared Nile Rat to eight other species of *Myomorpha* - lesser Egyptian jerboa *Jaculus jaculus*, Eurasian water vole *Arvicola amphibius*, golden hamster *Mesocricetus auratus*, white-footed mouse *Peromyscus leucopus*, Mongolian gerbil *Meriones unguiculatus*, house mouse *Mus musculus*, brown/Norway rat *Rattus norvegicus*, and human as an outgroup. To mitigate the effects of assembly quality and isoforms from different versions of assemblies, we re-annotated protein-coding genes of the 9 *Myomorpha* species by exonerate v2.4 (Slater and Birney 2005) using 20,426 human gene models that were generated by selecting the longest isoform and removing the pseudogenes.

After excluding genes lost in any taxa, a total of 19,628 orthologous genes remained for protein alignment. For detecting PSGs, we tested only candidates that passed a series of rigorous filters: 1) each gene had to map to the human gene with at least 70% coverage; 2) frameshift indels in coding sequences (CDS) were prohibited; 3) genes with premature stop codons were ruled out. A total of 7,492 high-quality orthologous genes remained.

The positive selection sites in Nile rat were detected by the branch-site model using PAML (v4.9j). Genes with an FDR-adjusted p-value less than 0.05 were treated as candidates for positive selection. To minimize effects of assembly and sequence alignment, we filtered positive selective sites by the following criteria: 1) the positive selective site was a gap in more than two species; 2) the PSG sites have more than two nonsynonymous substitution forms (ignoring the outgroup). We also performed a manual check for all individual PSGs to remove any other false-positives caused by low-quality alignments. This procedure detected 119 PSGs.

### Identification of diabetes-linked genes by text mining

We used four techniques to derive a set of genes associated with type 2 diabetes and with diet-induced diabetes. First we compiled an expert-curated gene-disease association database from standard resources, the Comparative Toxicogenomics Database (Davis et al. 2021) and PharmGKB (Thorn, Klein, and Altman 2013). The result gave 277 genes associated with type 2 diabetes, but none associated with diet-induced diabetes. Next, we employed Kinderminer, a simple text mining system developed to query ∼32 million PubMed abstracts to retrieve significantly associated target terms (e.g. genes) for an input key phrase (e.g. type 2 diabetes, diet-induced diabetes). KinderMiner retrieved 460 genes for type 2 diabetes and four genes for diet-induced diabetes. Third, we applied Serial KinderMiner (SKiM), a literature-based discovery system (LBD) that extends KinderMiner, querying the PubMed abstracts to find C terms (e.g. genes) for an input A term (e.g. type 2 diabetes, diet-induced diabetes) via some intermediate B terms (i.e. a list of phenotypes and symptoms). The set of B terms comprised only the top 50 phenotypes and symptoms significantly associated with type 2 diabetes or diet-induced diabetes. SKiM yielded 1,941 genes for type 2 diabetes and 2,254 genes for diet-induced diabetes. Restriction of the SKiM run to the top 50 phenotypes and symptoms ranked based on a prediction score is commonly practiced in other existing LBD systems such as LION LBD (Pyysalo et al. 2019) and BITOLA (Hristovski et al. 2006). In SKiM, the prediction score is calculated as a sum of negative logarithmic value of Fisher Exact Test (FET) p-value and sort ratio (i.e. the number of PubMed abstracts with A and B terms divided by the number of PubMed abstracts with B terms). Finally, we used a GWAS database (Buniello et al. 2019), which reported type 2 diabetes-associated SNPs in 1,482 genes.

We ranked the strength of association of each gene with diabetes as follows. An association reported in the gene-disease databases received a score of 3, an association reported by KinderMiner or the GWAS database received a score of 2, and SKiM the score of 1. If a gene was reported by more than one method, the scores were added up, so that the composite score ranged from 1 to 8.

## Data availability

### Primary genomic sequencing data

VGP GenomeArk: https://vgp.github.io/genomeark/Arvicanthis_niloticus/

### Transcriptomic sequencing data

We have deposited brain and testis Iso-seq data to the SRA:

- Brain: SRX8145073
- Testis: SRX8145073

### Genome assemblies and annotation at NCBI

Genome: https://www.ncbi.nlm.nih.gov/genome/?term=Arvicanthis+niloticus. Annotated reference genome assembly, genome browser, and other information are linked to from this page.

Reference, annotated genome assembly: mArvNil1.pat.X. This assembly contains paternal haplotype with maternal X chromosome. BioProject PRJNA632612; RefSeq Assembly GCF_011762505.1. Note: this version of the assembly does not contain the mitochondrion: NCBI was unable to include it for technical reasons. The maternal assembly is RefSeq Assembly GCA_011750645.1.

Nile rat genome annotation on NCBI FTP server: https://ftp.ncbi.nlm.nih.gov/genomes/all/annotation_releases/61156/100/

BioSample: SAMN12611849

Other genome assemblies in GenBank:

1. Principal pseudohaplotype: mArvNil1.pat.X. Contains paternal haplotype with maternal X chromosome. This assembly is identical to the RefSeq assembly but also contains the mitochondrial chromosome as a contig. BioProject PRJNA608735; Assembly GCA_011762505.1; mitochondrion CM022273
2. Paternal haplotype (uncurated): mArvNil1.pat. BioProject PRJNA561935; Assembly GCA_011762545.1
3. Maternal haplotype (uncurated): mArvNil1.mat. BioProject PRJNA561936; Assembly GCA_011750645.1. Note: this haplotype has not been curated. Therefore, some chromosomes remain split into 2 or more scaffolds.

## Supplementary datasets, code, and original high-resolution figures

Nile rat genome paper project on OSF: https://osf.io/j97kc/; DOI 10.17605/OSF.IO/J97KC Original high-resolution figures: https://osf.io/m3sjr/

TOGA annotation tracks for Nile rat genome, using human and mouse as the reference: https://genome.senckenberg.de//cgi-bin/hgTracks?db=HLarvNil1 (mArvNil1.pat.X assembly) https://genome.senckenberg.de//cgi-bin/hgTracks?db=HLarvNil1B (mArvNil1.mat assembly)

## Code availability

Data collection software was supplied by instrument vendors.

Free open source software was used for all types and stages of data analysis. See references in the Methods

Genome assembly and scaffolding: TrioCanu, Arrow, purge_dups, scaff10x, Bionano Solve, Salsa2, Longranger, Freebayes

Mitochondrial genome assembly: mitoVGP

Gene Ontology (GO) terms prediction: Phylo-PFP Segmental duplications workflow:

https://github.com/ChaissonLab/SegDupAnnotation/releases/tag/vNR

Segmental duplications workflow used windowmasker v1.0.0, RepeatMasker 4.1.1 with the parameter “-species rodentia”, SEDEF version 1.1-37-gd14abac-dirty with default parameters, CENSOR repeat masking server https://www.girinst.org/cgi-bin/censor/censor.cgi minimap2 was used for mapping gene models and Iso-seq transcripts to the genome Mutation rate analysis: GATK 4.0.7

Heterozygosity spectrum: Mummer v3.23, Assemblytics v1.2, SyRi v1.0

Branch-site test analysis: exonerate v2.4, PAML v4.9j

Text mining: KinderMiner, https://www.kinderminer.org/ and SKiM, https://skim.morgridge.org/ TOGA: https://github.com/hillerlab/TOGA

R:

R version 4.1.0 (2021-05-18)

Platform: x86_64-apple-darwin17.0 (64-bit)

Running under: macOS Big Sur 10.16

Matrix products: default

LAPACK: /Library/Frameworks/R.framework/Versions/4.1/Resources/lib/libRlapack.dylib

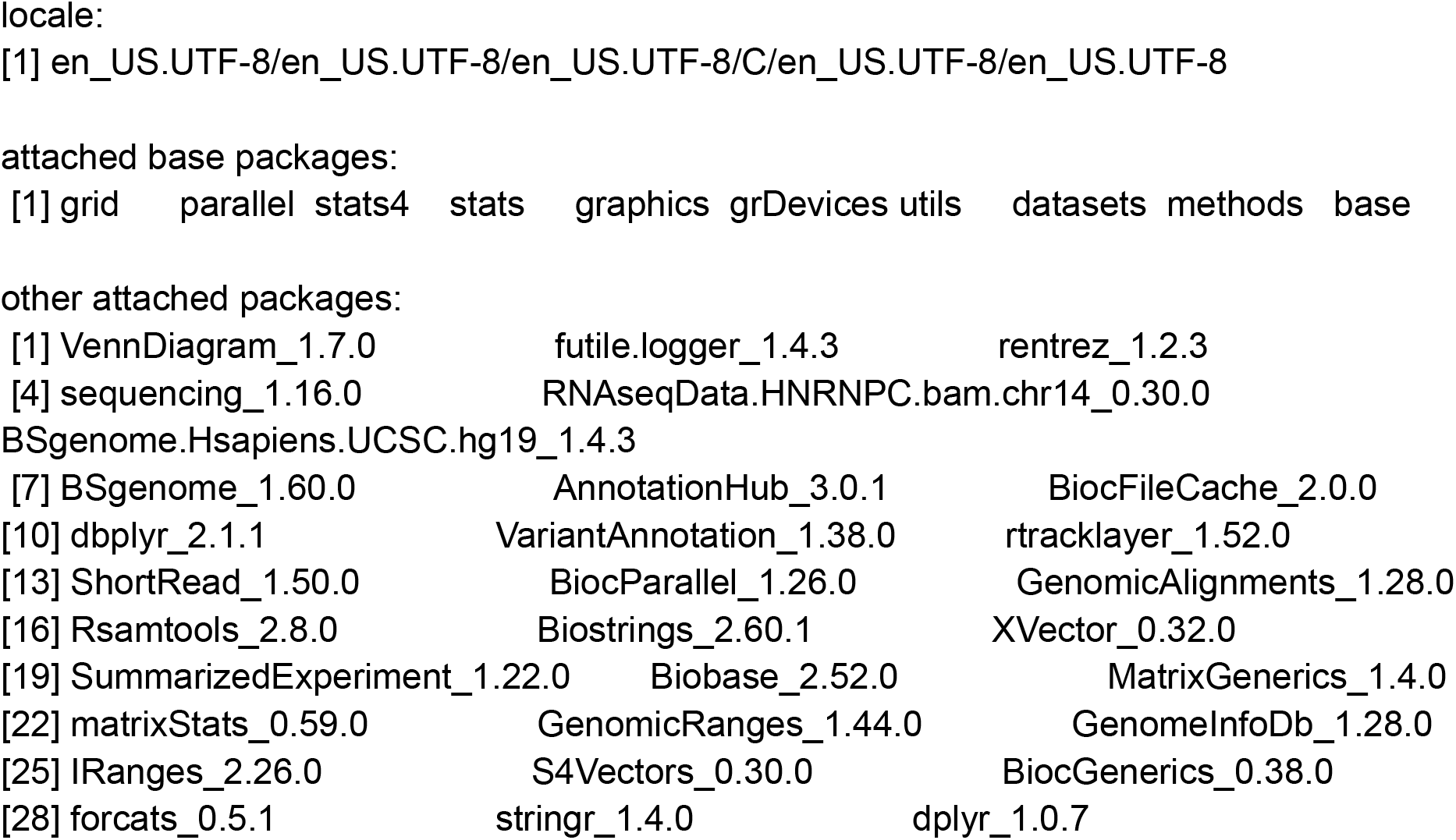
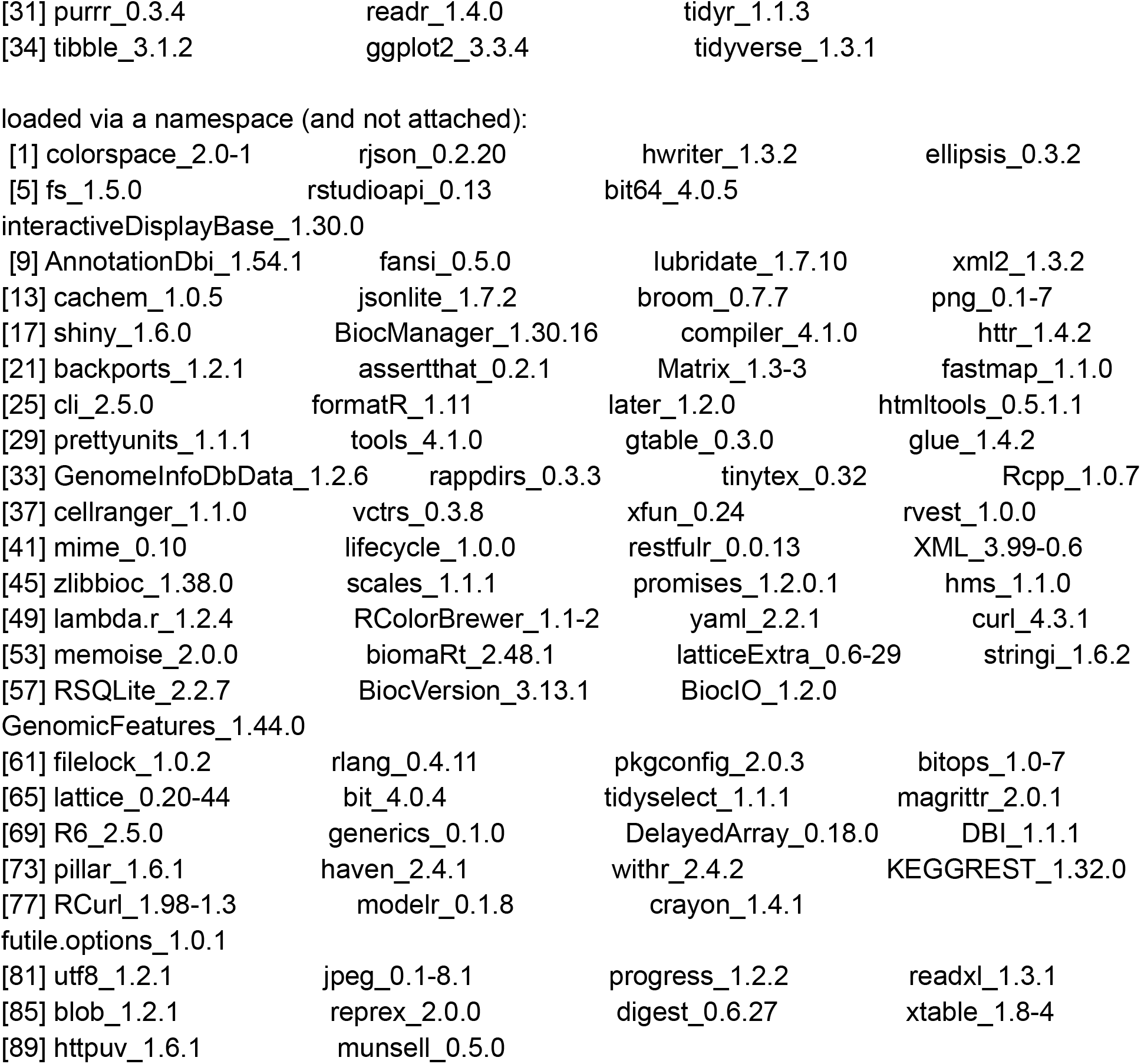

|  |  |  |
| --- | --- | --- |
| [1] VennDiagram_1.7.0 | futile.logger_1.4.3 | rentrez_1.2.3 |
| [4] sequencing_1.16.0 | RNAseqData.HNRNPC.bam.chr14_0.30.0 |  |
| BSgenome.Hsapiens.UCSC.hg19_1.4.3 |  |  |
| [7] BSgenome_1.60.0 | AnnotationHub_3.0.1 | BiocFileCache_2.0.0 |
| [10] dbplyr_2.1.1 | VariantAnnotation_1.38.0 | rtracklayer_1.52.0 |
| [13] ShortRead_1.50.0 | BiocParallel_1.26.0 | GenomicAlignments_1.28.0 |
| [16] Rsamtools_2.8.0 | Biostrings_2.60.1 | XVector_0.32.0 |
| [19] SummarizedExperiment_1.22.0 | Biobase_2.52.0 | MatrixGenerics_1.4.0 |
| [22] matrixStats_0.59.0 | GenomicRanges_1.44.0 | GenomeInfoDb_1.28.0 |
| [25] IRanges_2.26.0 | S4Vectors_0.30.0 | BiocGenerics_0.38.0 |
| [28] forcats_0.5.1 | stringr_1.4.0 | dplyr_1.0.7 |
| [31] purrr_0.3.4 | readr_1.4.0 | tidyr_1.1.3 |
| [34] tibble_3.1.2 | ggplot2_3.3.4 | tidyverse_1.3.1 |

|  |  |  |  |
| --- | --- | --- | --- |
| [1] colorspace_2.0-1 | rjson_0.2.20 | hwriter_1.3.2 | ellipsis_0.3.2 |
| [5] fs_1.5.0 | rstudioapi_0.13 | bit64_4.0.5 |  |
| interactiveDisplayBase_1.30.0 |  |  |  |
| [9] AnnotationDbi_1.54.1 | fansi_0.5.0 | lubridate_1.7.10 | xml2_1.3.2 |
| [13] cachem_1.0.5 | jsonlite_1.7.2 | broom_0.7.7 | png_0.1-7 |
| [17] shiny_1.6.0 | BiocManager_1.30.16 | compiler_4.1.0 | httr_1.4.2 |
| [21] backports_1.2.1 | assertthat_0.2.1 | Matrix_1.3-3 | fastmap_1.1.0 |
| [25] cli_2.5.0 | formatR_1.11 | later_1.2.0 | htmltools_0.5.1.1 |
| [29] prettyunits_1.1.1 | tools_4.1.0 | gtable_0.3.0 | glue_1.4.2 |
| [33] GenomeInfoDbData_1.2.6 | rappdirs_0.3.3 | tinytex_0.32 | Rcpp_1.0.7 |
| [37] cellranger_1.1.0 | vctrs_0.3.8 | xfun_0.24 | rvest_1.0.0 |
| [41] mime_0.10 | lifecycle_1.0.0 | restfulr_0.0.13 | XML_3.99-0.6 |
| [45] zlibbioc_1.38.0 | scales_1.1.1 | promises_1.2.0.1 | hms_1.1.0 |
| [49] lambda.r_1.2.4 | RColorBrewer_1.1-2 | yaml_2.2.1 | curl_4.3.1 |
| [53] memoise_2.0.0 | biomaRt_2.48.1 | latticeExtra_0.6-29 | stringi_1.6.2 |
| [57] RSQLite_2.2.7 | BiocVersion_3.13.1 | BiocIO_1.2.0 |  |
| GenomicFeatures_1.44.0 |  |  |  |
| [61] filelock_1.0.2 | rlang_0.4.11 | pkgconfig_2.0.3 | bitops_1.0-7 |
| [65] lattice_0.20-44 | bit_4.0.4 | tidyselect_1.1.1 | magrittr_2.0.1 |
| [69] R6_2.5.0 | generics_0.1.0 | DelayedArray_0.18.0 | DBI_1.1.1 |
| [73] pillar_1.6.1 | haven_2.4.1 | withr_2.4.2 | KEGGREST_1.32.0 |
| [77] RCurl_1.98-1.3 | modelr_0.1.8 | crayon_1.4.1 |  |
| futile.options_1.0.1 |  |  |  |
| [81] utf8_1.2.1 | jpeg_0.1-8.1 | progress_1.2.2 | readxl_1.3.1 |
| [85] blob_1.2.1 | reprex_2.0.0 | digest_0.6.27 | xtable_1.8-4 |
| [89] httpuv_1.6.1 | munsell_0.5.0 |  |  |

## Acknowledgments

This study was supported by The Garland Initiative for Vision funded by the William K. Bowes Jr. Foundation.

RS and JAT acknowledge a grant from Marv Conney.

MJPC is funded by NSF Grant Number 2046753.

MH acknowledges support from the LOEWE-Centre for Translational Biodiversity Genomics. (TBG) funded by the Hessen State Ministry of Higher Education, Research and the Arts (HMWK).

AR and AMP were supported by the Intramural Research Program of the National Human Genome Research Institute.

B Haase, J Mountcastle, O Fedrigo, and E.D. Jarvis’ contributions were supported by the Howard Hughes Medical Institute and the Rockefeller University.

Woori Kwak of C&K Genomics submitted data to NCBI.

Michael Collins set up software tools and the computing environment at Morgridge Institute for Research.

Francoise Thibaud-Nissen led the genome annotation effort at NCBI.

Amy Freitag helped edit the text.

We also thank Mark Springer, Ben-Yang Liao, David Thybert, Masa Roller, Cecile Ane, Noga Kronfeld-Schor, and Francoise Thibaud-Nissen for stimulating discussions and advice.

Alice Young and the NIH Intramural Sequencing Center (NISC) for assistance with sequencing.

## Author information

## Contributions

RS and JAT conceived of the project and provided guidance

HT, MJPC, CY, and YVB carried out most aspects of genome analysis and co-wrote the paper

HT provided Nile rat samples

JM, BH, and OF carried out genome sequencing and mapping assays AMP coordinated Illumina sequencing of the parents

GF, AR, and AMP assembled the genome AT, WC, and KH curated the genome

KR mined databases and literature for diabetes-linked genes

AJ and DK predicted GO terms and carried out gene set enrichment analyses

BK, CM, and MH projected human and mouse genes to the Nile rat genome

CY and GZ carried out heterozygosity spectrum and branch-site test analyses

JF provided phylogeny for the branch-site test analysis

LAB computed Nile rat mutation rate LY helped write the Introduction

SAS, PJ, and DOC provided guidance and suggestions and helped write the manuscript

YVB and EDJ coordinated collaborations and provided guidance

All of the authors read and approved the final manuscript

## Corresponding authors

Yury V. Bukhman, Mark J.P. Chaisson, Ron Stewart, and James A. Thomson

## Supplementary information

A supplement is included as a separate document Supplementary data on OSF: https://osf.io/j97kc/

## Bibliography

Bailey, Jeffrey A., and Evan E. Eichler. 2006. “Primate Segmental Duplications: Crucibles of Evolution, Diversity and Disease.” Nature Reviews. Genetics 7 (7): 552–64.

Bergeron, Lucie A., Søren Besenbacher, Tychele N. Turner, Cyril J. Versoza, Richard J. Wang, Alivia Lee Price, Ellie Armstrong, et al. 2021. “Mutationathon: Towards Standardization in Estimates of Pedigree-Based Germline Mutation Rates.” bioRxiv. https://doi.org/10.1101/2021.08.30.458162.

Bolsinger, Julia, Michelle Landstrom, Andrzej Pronczuk, Andrew Auerbach, and K. C. Hayes. 2017. “Low Glycemic Load Diets Protect against Metabolic Syndrome and Type 2 Diabetes Mellitus in the Male Nile Rat.” The Journal of Nutritional Biochemistry 42 (April): 134–48.

Buniello, Annalisa, Jacqueline A. L. MacArthur, Maria Cerezo, Laura W. Harris, James Hayhurst, Cinzia Malangone, Aoife McMahon, et al. 2019. “The NHGRI-EBI GWAS Catalog of Published Genome-Wide Association Studies, Targeted Arrays and Summary Statistics 2019.” Nucleic Acids Research 47 (D1): D1005–12.

Cao, Hongzhi, Honglong Wu, Ruibang Luo, Shujia Huang, Yuhui Sun, Xin Tong, Yinlong Xie, et al. 2015. “De Novo Assembly of a Haplotype-Resolved Human Genome.” Nature Biotechnology 33 (6): 617–22.

Cederroth, Christopher R., Urs Albrecht, Joseph Bass, Steven A. Brown, Jonas Dyhrfjeld-Johnsen, Frederic Gachon, Carla B. Green, et al. 2019. “Medicine in the Fourth Dimension.” Cell Metabolism 30 (2): 238–50.

Chaabo, Fadi, Andrzej Pronczuk, Ekaterina Maslova, and Kc Hayes. 2010. “Nutritional Correlates and Dynamics of Diabetes in the Nile Rat (Arvicanthis Niloticus): A Novel Model for Diet-Induced Type 2 Diabetes and the Metabolic Syndrome.” Nutrition & Metabolism 7 (April): 29.

Chen, Ji, Cassandra N. Spracklen, Gaëlle Marenne, Arushi Varshney, Laura J. Corbin, Jian ‘an Luan, Sara M. Willems, et al. 2021. “The Trans-Ancestral Genomic Architecture of Glycemic Traits.” Nature Genetics 53 (6): 840–60.

Cheung, Joseph, Michael D. Wilson, Junjun Zhang, Razi Khaja, Jeffrey R. MacDonald, Henry H. Q. Heng, Ben F. Koop, and Stephen W. Scherer. 2003. “Recent Segmental and Gene Duplications in the Mouse Genome.” Genome Biology 4 (8): R47.

Collin, Gayle B., Jan D. Marshall, Akihiro Ikeda, W. Venus So, Isabelle Russell-Eggitt, Pietro Maffei, Sebastian Beck, et al. 2002. “Mutations in ALMS1 Cause Obesity, Type 2 Diabetes and Neurosensory Degeneration in Alström Syndrome.” Nature Genetics 31 (1): 74–78.

Consortium, Mouse Genome Sequencing, and Mouse Genome Sequencing Consortium. 2002. “Initial Sequencing and Comparative Analysis of the Mouse Genome.” Nature. https://doi.org/10.1038/nature01262.

Consortium, Rat Genome Sequencing Project, and Rat Genome Sequencing Project Consortium. 2004. “Genome Sequence of the Brown Norway Rat Yields Insights into Mammalian Evolution.” Nature. https://doi.org/10.1038/nature02426.

Cooke, Amy, Thomas Schwarzl, Ina Huppertz, Gertjan Kramer, Panagiotis Mantas, Anne-Marie Alleaume, Wolfgang Huber, Jeroen Krijgsveld, and Matthias W. Hentze. 2019. “The RNA-Binding Protein YBX3 Controls Amino Acid Levels by Regulating SLC mRNA Abundance.” Cell Reports 27 (11): 3097–3106.e5.

Davis, Allan Peter, Cynthia J. Grondin, Robin J. Johnson, Daniela Sciaky, Jolene Wiegers, Thomas C. Wiegers, and Carolyn J. Mattingly. 2021. “Comparative Toxicogenomics Database (CTD): Update 2021.” Nucleic Acids Research 49 (D1): D1138–43.

Diabetes Genetics Initiative of Broad Institute of Harvard and MIT, Lund University, and Novartis Institutes of BioMedical Research, Richa Saxena, Benjamin F. Voight, Valeriya Lyssenko, Noël P. Burtt, Paul I. W. de Bakker, Hong Chen, et al. 2007. “Genome-Wide Association Analysis Identifies Loci for Type 2 Diabetes and Triglyceride Levels.” Science 316 (5829): 1331–36.

Falchi, Mario, Julia Sarah El-Sayed Moustafa, Petros Takousis, Francesco Pesce, Amélie Bonnefond, Johanna C. Andersson-Assarsson, Peter H. Sudmant, et al. 2014. “Low Copy Number of the Salivary Amylase Gene Predisposes to Obesity.” Nature Genetics 46 (5): 492–97.

Flanagan, Sarah E., Ann-Marie Patch, and Sian Ellard. 2010. “Using SIFT and PolyPhen to Predict Loss-of-Function and Gain-of-Function Mutations.” Genetic Testing and Molecular Biomarkers 14 (4): 533–37.

Formenti, Giulio, Arang Rhie, Jennifer Balacco, Bettina Haase, Jacquelyn Mountcastle, Olivier Fedrigo, Samara Brown, et al. 2021. “Complete Vertebrate Mitogenomes Reveal Widespread Repeats and Gene Duplications.” Genome Biology 22 (1): 120.

Foti, Daniela, Eusebio Chiefari, Monica Fedele, Rodolfo Iuliano, Leonardo Brunetti, Francesco Paonessa, Guidalberto Manfioletti, et al. 2005. “Lack of the Architectural Factor HMGA1 Causes Insulin Resistance and Diabetes in Humans and Mice.” Nature Medicine 11 (7): 765–73.

Frühbeck, Gema, Blanca Fernández-Quintana, Mirla Paniagua, Ana Wenting Hernández-Pardos, Víctor Valentí, Rafael Moncada, Victoria Catalán, et al. 2020. “FNDC4, a Novel Adipokine That Reduces Lipogenesis and Promotes Fat Browning in Human Visceral Adipocytes.” Metabolism: Clinical and Experimental 108 (July): 154261.

Gaillard, Frédéric, Stephan Bonfield, Gregory S. Gilmour, Sharee Kuny, Silvina C. Mema, Brent T. Martin, Laura Smale, Nathan Crowder, William K. Stell, and Yves Sauvé. 2008. “Retinal Anatomy and Visual Performance in a Diurnal Cone-Rich Laboratory Rodent, the Nile Grass Rat (Arvicanthis Niloticus).” The Journal of Comparative Neurology. https://doi.org/10.1002/cne.21798.

Gaillard, Frédéric, Harvey J. Karten, and Yves Sauvé. 2013. “Retinorecipient Areas in the Diurnal Murine Rodent Arvicanthis Niloticus: A Disproportionally Large Superior Colliculus.” The Journal of Comparative Neurology 521 (8): Spc1.

Goel, Manish, Hequan Sun, Wen-Biao Jiao, and Korbinian Schneeberger. 2019. “SyRI: Finding Genomic Rearrangements and Local Sequence Differences from Whole-Genome Assemblies.” Genome Biology 20 (1): 277.

Gordon, David, John Huddleston, Mark J. P. Chaisson, Christopher M. Hill, Zev N. Kronenberg, Katherine M. Munson, Maika Malig, et al. 2016. “Long-Read Sequence Assembly of the Gorilla Genome.” Science 352 (6281): aae0344.

Gotfryd, Kamil, Andreia Filipa Mósca, Julie Winkel Missel, Sigurd Friis Truelsen, Kaituo Wang, Mariana Spulber, Simon Krabbe, et al. 2018. “Human Adipose Glycerol Flux Is Regulated by a pH Gate in AQP10.” Nature Communications 9 (1): 4749.

Guan, Dengfeng, Shane A. McCarthy, Jonathan Wood, Kerstin Howe, Yadong Wang, and Richard Durbin. 2020. “Identifying and Removing Haplotypic Duplication in Primary Genome Assemblies.” Bioinformatics 36 (9): 2896–98.

Gudmundsson, Sanna, Konrad J. Karczewski, Laurent C. Francioli, Grace Tiao, Beryl B. Cummings, Jessica Alföldi, Qingbo Wang, et al. 2021. “Addendum: The Mutational Constraint Spectrum Quantified from Variation in 141,456 Humans.” Nature 597 (7874): E3–4.

Hernández-Saavedra, Diego, and Kristin I. Stanford. 2019. “The Regulation of Lipokines by Environmental Factors.” Nutrients 11 (10). https://doi.org/10.3390/nu11102422.

Heymann, Anthony D., Yossi Cohen, and Gabriel Chodick. 2012. “Glucose-6-Phosphate Dehydrogenase Deficiency and Type 2 Diabetes.” Diabetes Care 35 (8): e58.

Howe, Kerstin, William Chow, Joanna Collins, Sarah Pelan, Damon-Lee Pointon, Ying Sims, James Torrance, Alan Tracey, and Jonathan Wood. 2020. “Significantly Improving the Quality of Genome Assemblies through Curation.” bioRxiv. https://doi.org/10.1101/2020.08.12.247734.

Hristovski, Dimitar, Carol Friedman, Thomas C. Rindflesch, and Borut Peterlin. 2006. “Exploiting Semantic Relations for Literature-Based Discovery.” AMIA … Annual Symposium Proceedings / AMIA Symposium. AMIA Symposium, 349–53.

Hunt, Mary C., and Stefan E. H. Alexson. 2008. “Novel Functions of Acyl-CoA Thioesterases and Acyltransferases as Auxiliary Enzymes in Peroxisomal Lipid Metabolism.” Progress in Lipid Research 47 (6): 405–21.

Jain, Aashish, and Daisuke Kihara. 2019. “Phylo-PFP: Improved Automated Protein Function Prediction Using Phylogenetic Distance of Distantly Related Sequences.” Bioinformatics 35 (5): 753–59.

Kalsbeek, Andries, Linda A. W. Verhagen, Ingrid Schalij, Ewout Foppen, Michel Saboureau, Béatrice Bothorel, Ruud M. Buijs, and Paul Pévet. 2008. “Opposite Actions of Hypothalamic Vasopressin on Circadian Corticosterone Rhythm in Nocturnal versus Diurnal Species.” The European Journal of Neuroscience 27 (4): 818–27.

Kirilenko, Bogdan M., Lee R. Hagey, Stephen Barnes, Charles N. Falany, and Michael Hiller. 2019. “Evolutionary Analysis of Bile Acid-Conjugating Enzymes Reveals a Complex Duplication and Reciprocal Loss History.” Genome Biology and Evolution 11 (11): 3256–68.

Kirilenko, Bogdan M., Chetan Munegowda, Ekaterina Osipova, David Jebb, Virag Sharma, Moritz Blumer, Ariadna Morales, et al. Submitted. “TOGA Integrates Gene Annotation with Orthology Inference at Scale.”

Koren, Sergey, Arang Rhie, Brian P. Walenz, Alexander T. Dilthey, Derek M. Bickhart, Sarah B. Kingan, Stefan Hiendleder, John L. Williams, Timothy P. L. Smith, and Adam M. Phillippy. 2018. “De Novo Assembly of Haplotype-Resolved Genomes with Trio Binning.” Nature Biotechnology, October. https://doi.org/10.1038/nbt.4277.

Kottaisamy, Chidhambara Priya Dharshini, Divya S. Raj, V. Prasanth Kumar, and Umamaheswari Sankaran. 2021. “Experimental Animal Models for Diabetes and Its Related Complications-a Review.” Laboratory Animal Research 37 (1): 23.

Kriventseva, Evgenia V., Dmitry Kuznetsov, Fredrik Tegenfeldt, Mosè Manni, Renata Dias Felipe, A. Simão, and Evgeny M. Zdobnov. 2019. “OrthoDB v10: Sampling the Diversity of Animal, Plant, Fungal, Protist, Bacterial and Viral Genomes for Evolutionary and Functional Annotations of Orthologs.” Nucleic Acids Research 47 (D1): D807–11.

Kuusisto, Finn, Daniel Ng, John Steill, Ian Ross, Miron Livny, James Thomson, David Page, and Ron Stewart. 2020. “KinderMiner Web: A Simple Web Tool for Ranking Pairwise Associations in Biomedical Applications.” F1000Research 9 (832): 832.

Labay, Valentina, Tal Raz, Dana Baron, Hanna Mandel, Hawys Williams, Timothy Barrett, Raymonde Szargel, et al. 1999. “Mutations in SLC19A2 Cause Thiamine-Responsive Megaloblastic Anaemia Associated with Diabetes Mellitus and Deafness.” Nature Genetics 22 (3): 300–304.

Langel, Jennifer, Tomoko Ikeno, Lily Yan, Antonio A. Nunez, and Laura Smale. 2018. “Distributions of GABAergic and Glutamatergic Neurons in the Brains of a Diurnal and Nocturnal Rodent.” Brain Research 1700 (December): 152–59.

Lee, Wonjae, Subin Yun, Geum Hee Choi, and Tae Woo Jung. 2018. “Fibronectin Type III Domain Containing 4 Attenuates Hyperlipidemia-Induced Insulin Resistance via Suppression of Inflammation and ER Stress through HO-1 Expression in Adipocytes.” Biochemical and Biophysical Research Communications 502 (1): 129–36.

Lilue, Jingtao, Anthony G. Doran, Ian T. Fiddes, Monica Abrudan, Joel Armstrong, Ruth Bennett, William Chow, et al. 2018. “Sixteen Diverse Laboratory Mouse Reference Genomes Define Strain-Specific Haplotypes and Novel Functional Loci.” Nature Genetics 50 (11): 1574–83.

Long, Anthony D., James Baldwin-Brown, Yuan Tao, Vanessa J. Cook, Gabriela Balderrama-Gutierrez, Russell Corbett-Detig, Ali Mortazavi, and Alan G. Barbour. 2019. “The Genome of Peromyscus Leucopus, Natural Host for Lyme Disease and Other Emerging Infections.” Science Advances 5 (7): eaaw6441.

Malik, Vasanti S., Barry M. Popkin, George A. Bray, Jean-Pierre Després, and Frank B. Hu. 2010. “Sugar-Sweetened Beverages, Obesity, Type 2 Diabetes Mellitus, and Cardiovascular Disease Risk.” Circulation 121 (11): 1356–64.

McElhinny, Teresa L., Laura Smale, and Kay E. Holekamp. 1997. “Patterns of Body Temperature, Activity, and Reproductive Behavior in a Tropical Murid Rodent, Arvicanthis Niloticus.” Physiology & Behavior. https://doi.org/10.1016/s0031-9384(97)00146-7.

Morgulis, Aleksandr, E. Michael Gertz Alejandro, A. Schäffer, and Richa Agarwala. 2006. “WindowMasker: Window-Based Masker for Sequenced Genomes.” Bioinformatics 22 (2): 134–41.

Morinaga, Tomonori, Masamichi Nakakoshi, Atsushi Hirao, Masashi Imai, and Kenichi Ishibashi. 2002. “Mouse Aquaporin 10 Gene (AQP10) Is a Pseudogene.” Biochemical and Biophysical Research Communications 294 (3): 630–34.

Nattestad, Maria, and Michael C. Schatz. 2016. “Assemblytics: A Web Analytics Tool for the Detection of Variants from an Assembly.” Bioinformatics 32 (19): 3021–23.

Numanagic, Ibrahim, Alim S. Gökkaya, Lillian Zhang, Bonnie Berger, Can Alkan, and Faraz Hach. 2018. “Fast Characterization of Segmental Duplications in Genome Assemblies.” Bioinformatics 34 (17): i706–14.

O’Leary Nuala A., Mathew W. Wright, J. Rodney Brister, Stacy Ciufo, Diana Haddad, Rich McVeigh, Bhanu Rajput, et al. 2016. “Reference Sequence (RefSeq) Database at NCBI: Current Status, Taxonomic Expansion, and Functional Annotation.” Nucleic Acids Research 44 (D1): D733–45.

Parsons, J. D. 1995. “Miropeats: Graphical DNA Sequence Comparisons.” Computer Applications in the Biosciences: CABIOS 11 (6): 615–19.

Pautsch, Alexander, Nadja Stadler, Adelheid Löhle, Wolfgang Rist, Adina Berg, Lucia Glocker, Herbert Nar, et al. 2013. “Crystal Structure of Glucokinase Regulatory Protein.” Biochemistry 52 (20): 3523–31.

Pedersen, Helle Krogh, Valborg Gudmundsdottir, and Søren Brunak. 2017. “Pancreatic Islet Protein Complexes and Their Dysregulation in Type 2 Diabetes.” Frontiers in Genetics 8 (April): 43.

Perry, George H., Nathaniel J. Dominy, Katrina G. Claw, Arthur S. Lee, Heike Fiegler, Richard Redon, John Werner, et al. 2007. “Diet and the Evolution of Human Amylase Gene Copy Number Variation.” Nature Genetics 39 (10): 1256–60.

Plesner, Annette, Peter Liston, Rusung Tan, Robert G. Korneluk, and C. Bruce Verchere. 2005. “The X-Linked Inhibitor of Apoptosis Protein Enhances Survival of Murine Islet Allografts.” Diabetes 54 (9): 2533–40.

Pyysalo, Sampo, Simon Baker, Imran Ali, Stefan Haselwimmer, Tejas Shah, Andrew Young, Yufan Guo, et al. 2019. “LION LBD: A Literature-Based Discovery System for Cancer Biology.” Bioinformatics 35 (9): 1553–61.

Raimondo, Anne, Matthew G. Rees, and Anna L. Gloyn. 2015. “Glucokinase Regulatory Protein: Complexity at the Crossroads of Triglyceride and Glucose Metabolism.” Current Opinion in Lipidology 26 (2): 88–95.

Raja, Kalpana, John Steill, Ian Ross, Lam C. Tsoi, Finn Kuusisto, Zijian Ni, Miron Livny, James Thomson, and Ron Stewart. 2020. “SKiM - A Generalized Literature-Based Discovery System for Uncovering Novel Biomedical Knowledge from PubMed.” bioRxiv. https://doi.org/10.1101/2020.10.16.343012.

Rhie, Arang, Shane A. McCarthy, Olivier Fedrigo, Joana Damas, Giulio Formenti, Sergey Koren, Marcela Uliano-Silva, et al. 2021. “Towards Complete and Error-Free Genome Assemblies of All Vertebrate Species.” Nature 592 (7856): 737–46.

Sanghera, Dharambir K., Ruth Hopkins, Megan W. Malone-Perez, Cynthia Bejar, Chengcheng Tan, Huda Mussa, Paul Whitby, et al. 2019. “Targeted Sequencing of Candidate Genes of Dyslipidemia in Punjabi Sikhs: Population-Specific Rare Variants in GCKR Promote Ectopic Fat Deposition.” PloS One 14 (8): e0211661.

Sarsani, Vishal Kumar, Narayanan Raghupathy, Ian T. Fiddes, Joel Armstrong, Francoise Thibaud-Nissen, Oraya Zinder, Mohan Bolisetty, et al. 2019. “The Genome of C57BL/6J ‘Eve’, the Mother of the Laboratory Mouse Genome Reference Strain.” G3 Genes|Genomes|Genetics 9 (6): 1795–1805.

Shimoyama, Mary, Jeff De Pons, G. Thomas Hayman, Stanley J. F. Laulederkind, Weisong Liu, Rajni Nigam, Victoria Petri, et al. 2015. “The Rat Genome Database 2015: Genomic, Phenotypic and Environmental Variations and Disease.” Nucleic Acids Research 43 (Database issue): D743–50.

Sibly, Richard M., and James H. Brown. 2007. “Effects of Body Size and Lifestyle on Evolution of Mammal Life Histories.” Proceedings of the National Academy of Sciences of the United States of America 104 (45): 17707–12.

Slater, Guy St C., and Ewan Birney. 2005. “Automated Generation of Heuristics for Biological Sequence Comparison.” BMC Bioinformatics 6 (February): 31.

Smit, Arian F. A., Robert Hubley, and P. Green. 1996. “RepeatMasker.”

Sparsø, T., G. Andersen, T. Nielsen, K. S. Burgdorf, A. P. Gjesing, A. L. Nielsen, A. Albrechtsen, et al. 2008. “The GCKR rs780094 Polymorphism Is Associated with Elevated Fasting Serum Triacylglycerol, Reduced Fasting and OGTT-Related Insulinaemia, and Reduced Risk of Type 2 Diabetes.” Diabetologia 51 (1): 70–75.

Sulak, Michael, Lindsey Fong, Katelyn Mika, Sravanthi Chigurupati, Lisa Yon, Nigel P. Mongan, Richard D. Emes, and Vincent J. Lynch. 2016. “TP53 Copy Number Expansion Is Associated with the Evolution of Increased Body Size and an Enhanced DNA Damage Response in Elephants.” eLife 5 (September). https://doi.org/10.7554/eLife.11994.

Sullivan, S. L., M. C. Adamson, K. J. Ressler, C. A. Kozak, and L. B. Buck. 1996. “The Chromosomal Distribution of Mouse Odorant Receptor Genes.” Proceedings of the National Academy of Sciences of the United States of America 93 (2): 884–88.

Thibaud-Nissen, Françoise, Alexander Souvorov, Terence Murphy, Michael DiCuccio, and Paul Kitts. 2013. Eukaryotic Genome Annotation Pipeline. National Center for Biotechnology Information (US).

Thorn, Caroline F., Teri E. Klein, and Russ B. Altman. 2013. “PharmGKB: The Pharmacogenomics Knowledge Base.” Methods in Molecular Biology 1015: 311–20.

Thybert, David, Maša Roller, Fábio C. P. Navarro, Ian Fiddes, Ian Streeter, Christine Feig, David Martin-Galvez, et al. 2018. “Repeat Associated Mechanisms of Genome Evolution and Function Revealed by the and Genomes.” Genome Research 28 (4): 448–59.

Toh, Huishi, Alexander Smolentsev, Rachel V. Bozadjian, Patrick W. Keeley, Madison D. Lockwood, Ryan Sadjadi, Dennis O. Clegg, et al. 2019. “Vascular Changes in Diabetic Retinopathy-a Longitudinal Study in the Nile Rat.” Laboratory Investigation; a Journal of Technical Methods and Pathology 99 (10): 1547–60.

Toh, Huishi, James A. Thomson, and Peng Jiang. 2020. “Maternal High-Fiber Diet Protects Offspring against Type 2 Diabetes.” Nutrients 13 (1). https://doi.org/10.3390/nu13010094.

Tsuboi, Ayaka, Satomi Minato, Megumu Yano, Mika Takeuchi, Kaori Kitaoka, Miki Kurata, Gen Yoshino, Bin Wu, Tsutomu Kazumi, and Keisuke Fukuo. 2018. “Association of Serum Orosomucoid with 30-Min Plasma Glucose and Glucose Excursion during Oral Glucose Tolerance Tests in Non-Obese Young Japanese Women.” BMJ Open Diabetes Research & Care 6 (1): e000508.

Vaxillaire, Martine, Christine Cavalcanti-Proença, Aurélie Dechaume, Jean Tichet, Michel Marre, Beverley Balkau, Philippe Froguel, and DESIR Study Group. 2008. “The Common P446L Polymorphism in GCKR Inversely Modulates Fasting Glucose and Triglyceride Levels and Reduces Type 2 Diabetes Risk in the DESIR Prospective General French Population.” Diabetes 57 (8): 2253–57.

Vollger, Mitchell R., Philip C. Dishuck, Melanie Sorensen, Annemarie E. Welch, Vy Dang, Max L. Dougherty, Tina A. Graves-Lindsay, Richard K. Wilson, Mark J. P. Chaisson, and Evan E. Eichler. 2019. “Long-Read Sequence and Assembly of Segmental Duplications.” Nature Methods 16 (1): 88–94.

Volobouev, V. T., J. F. Ducroz, V. M. Aniskin, J. Britton-Davidian, R. Castiglia, G. Dobigny, L. Granjon, et al. 2002. “Chromosomal Characterization of Arvicanthis Species (Rodentia, Murinae) from Western and Central Africa: Implications for Taxonomy.” Cytogenetic and Genome Research 96 (1-4): 250–60.

Wang, Bo, Elizabeth Tseng, Michael Regulski, Tyson A. Clark, Ting Hon, Yinping Jiao, Zhenyuan Lu, Andrew Olson, Joshua C. Stein, and Doreen Ware. 2016. “Unveiling the Complexity of the Maize Transcriptome by Single-Molecule Long-Read Sequencing.” Nature Communications. https://doi.org/10.1038/ncomms11708.

White, Phillip J., Robert W. McGarrah, Mark A. Herman, James R. Bain, Svati H. Shah, and Christopher B. Newgard. 2021. “Insulin Action, Type 2 Diabetes, and Branched-Chain Amino Acids: A Two-Way Street.” Molecular Metabolism 52 (October): 101261.

Yang, Chentao, Yang Zhou, Stephanie Marcus, Giulio Formenti, Lucie A. Bergeron, Zhenzhen Song, Xupeng Bi, et al. 2021. “Evolutionary and Biomedical Insights from a Marmoset Diploid Genome Assembly.” Nature 594 (7862): 227–33.

Yang, Kaiyuan, Jonathan Gotzmann, Sharee Kuny, Hui Huang, Yves Sauvé, and Catherine B. Chan. 2016. “Five Stages of Progressive β-Cell Dysfunction in the Laboratory Nile Rat Model of Type 2 Diabetes.” The Journal of Endocrinology 229 (3): 343–56.

Yan, Lily, Laura Smale, and Antonio A. Nunez. 2020. “Circadian and Photic Modulation of Daily Rhythms in Diurnal Mammals.” The European Journal of Neuroscience 51 (1): 551–66.

Zahedi, Asiyeh Sadat, Mahdi Akbarzadeh, Bahareh Sedaghati-Khayat, Atefeh Seyedhamzehzadeh, and Maryam S. Daneshpour. 2021. “GCKR Common Functional Polymorphisms Are Associated with Metabolic Syndrome and Its Components: A 10-Year Retrospective Cohort Study in Iranian Adults.” Diabetology & Metabolic Syndrome 13 (1): 20.

