## Supplement for "A haplotype-resolved genome assembly of the Nile rat facilitates exploration of the genetic basis of diabetes"

Genes linked to type 2 diabetes

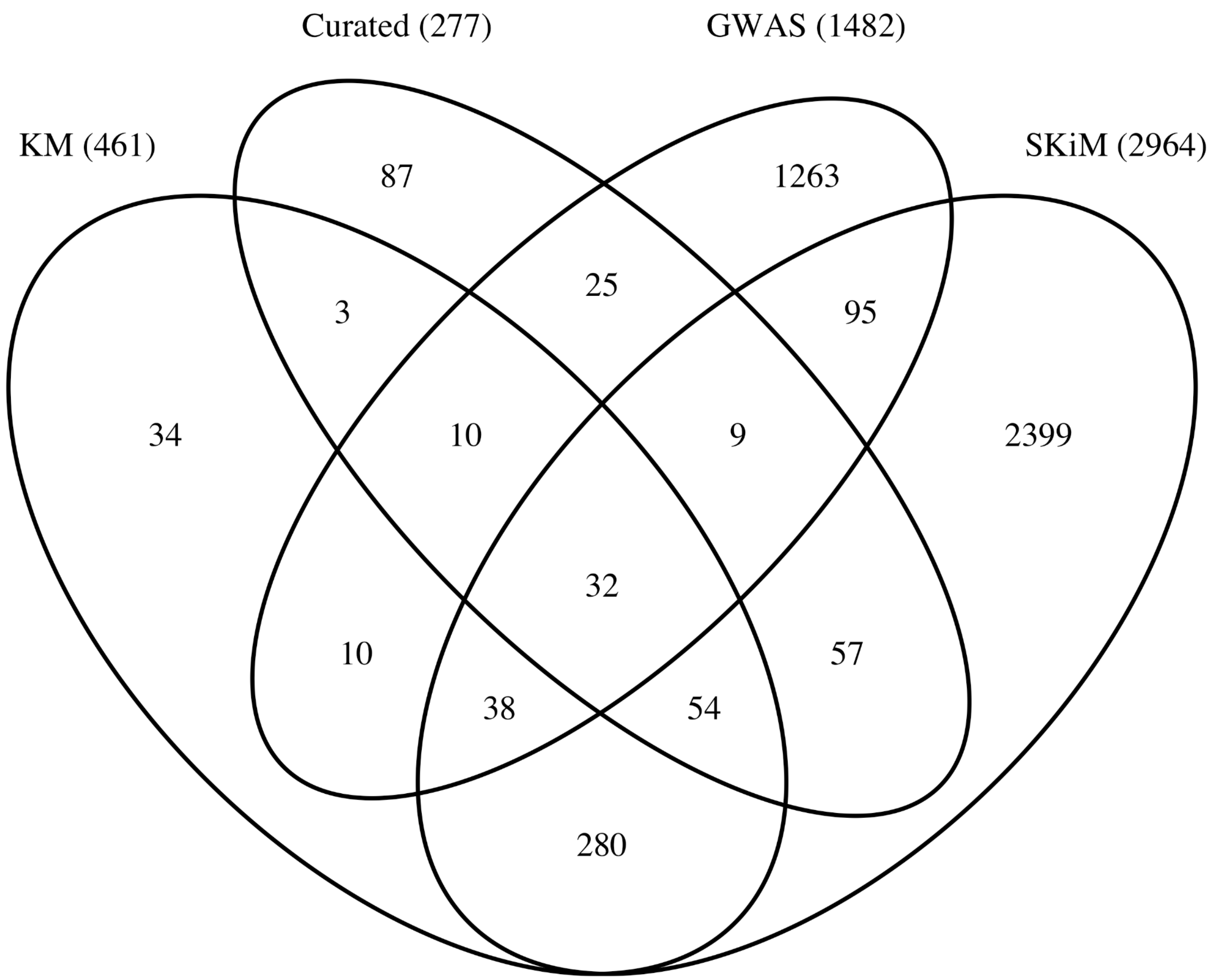


**Supplementary Figure 1. Venn diagram of gene lists linked to type 2 diabetes by different types of evidence.** KM = KinderMiner, a text-mining tool. SKiM - Serial KinderMiner, a literature-based discovery system.

### Heterozygosity spectrum of the Nile rat

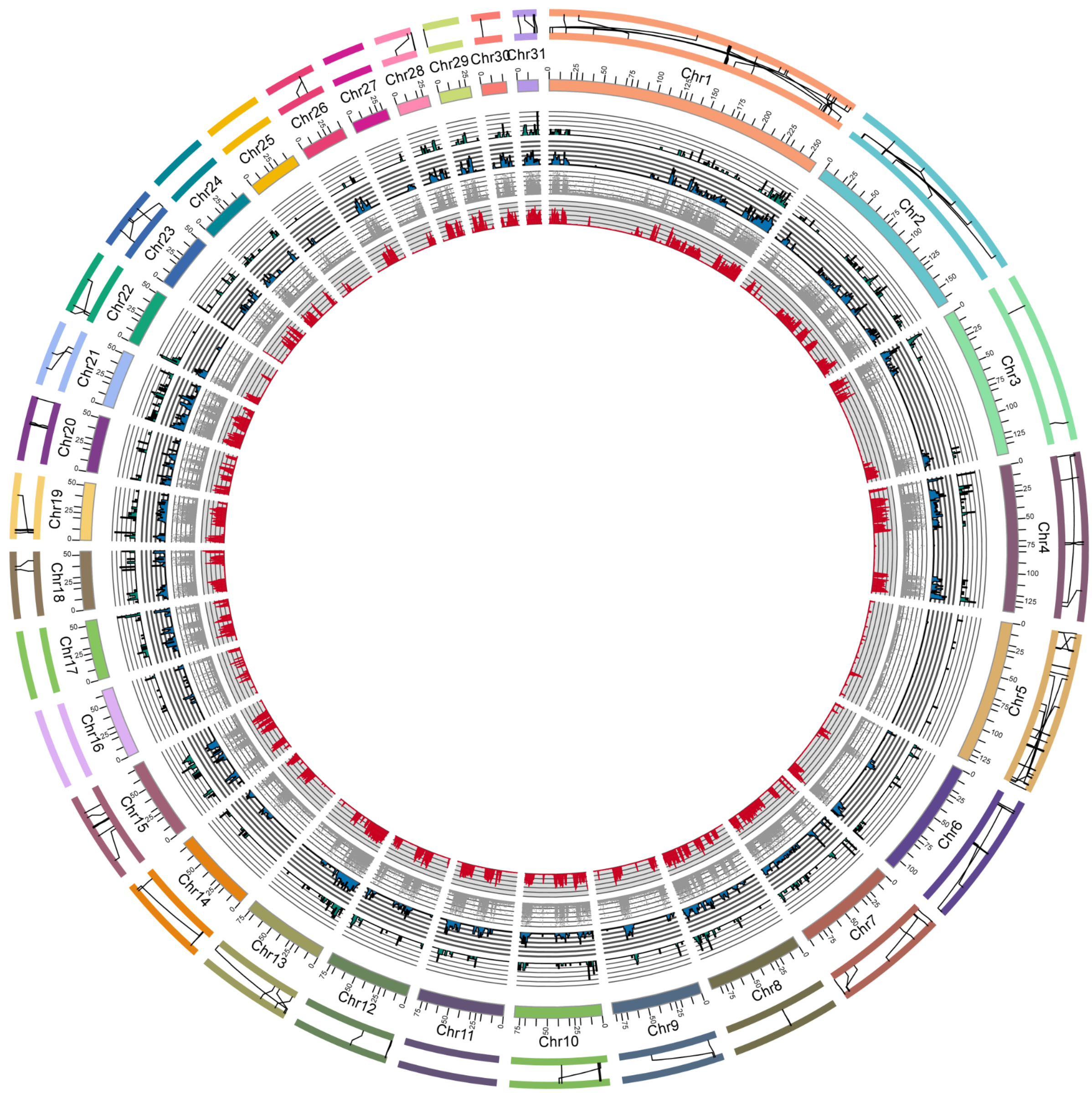


**Supplementary Figure 2. Heterozygosity inferred by comparing the paternal and maternal scaffolded contigs, shown on the paternal scaffolds.** From inner ring, the plots show density of heterozygous SNV, indels up to 50 bases, and finally insertion and deletion structural variants ( ≥ 50 bases). The outer rim shows translocations.

### Length distributions of structural variants

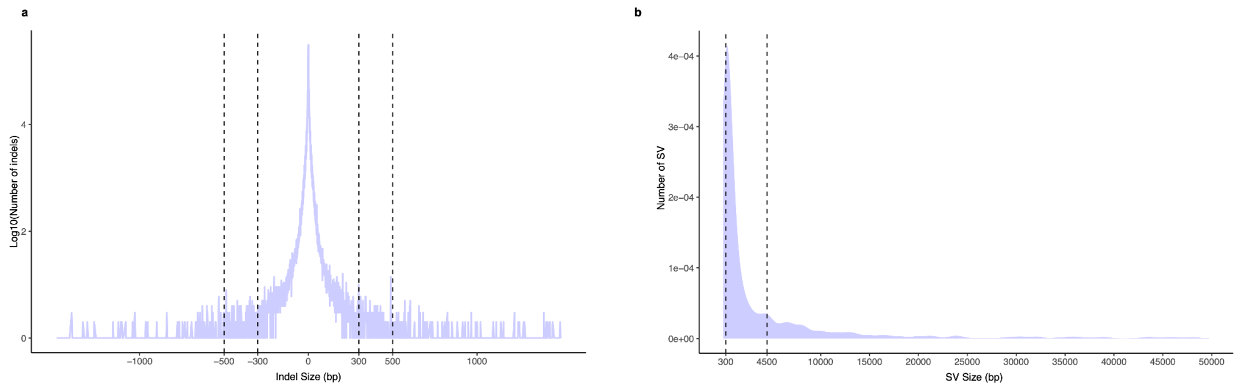


**Supplementary Figure 3. Length distributions of structural variants. A.** Indels. **B**. Other structural variants.

### Functional classification of duplicated genes.

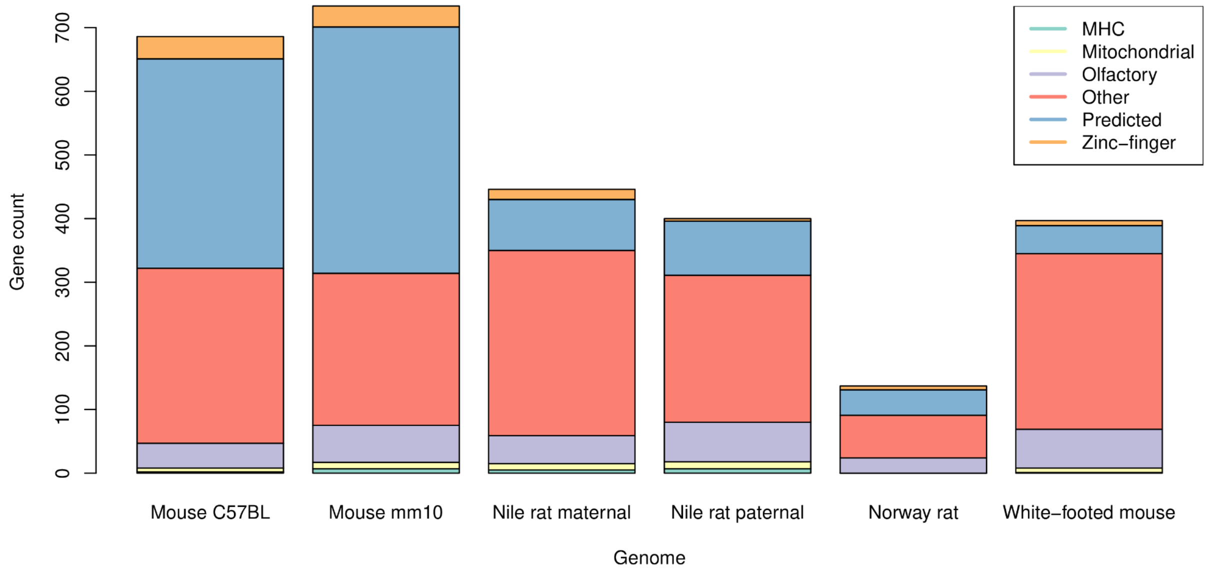


**Supplementary Figure 4. Functional classification of duplicated genes.** Genes are classified by name according to large categories of gene function, immune (MHC), Mitochondrial, Olfactory, Predicted, and Zinc-finger. The remaining category (Other) represents multi-copy genes not part of large gene families.

### Gckr

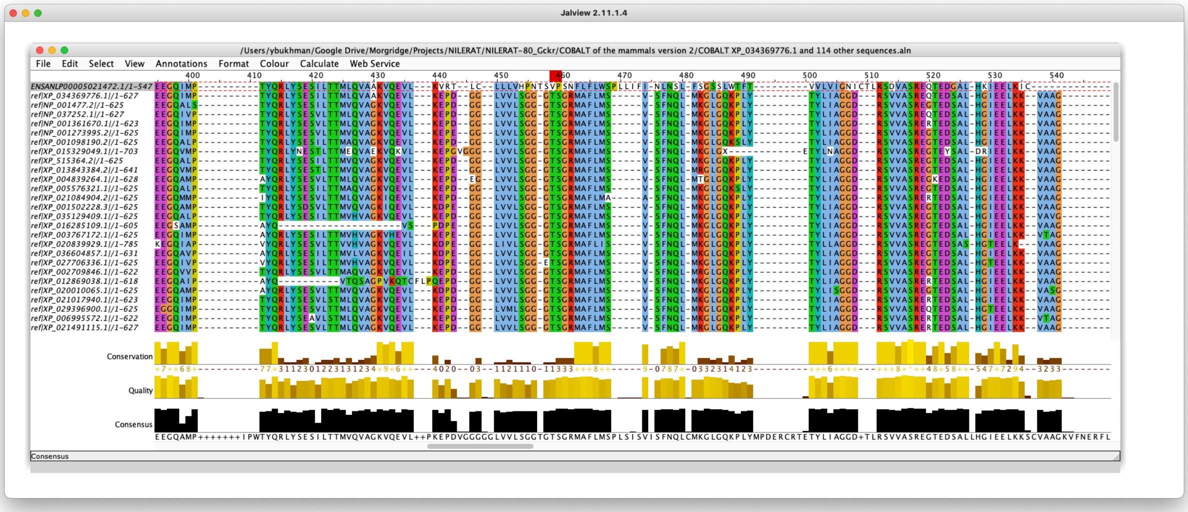


**Supplementary Figure 5.** A portion of a multiple alignment of the two Nile rat Gckr proteins to 113 mammalian orthologs displayed in JalView. The second copy of Nile rat Gckr is in the top row, canonical Nile rat Gckr in the second row, followed by human, rat, mouse, and other mammals. Positions 459 and 460 of the alignment correspond to T109 and S110 of the human protein. These constitute a part of the F1P binding site, with T109 involved in polar contacts with hydroxyl substituents of fructopyranose and S110 forming a hydrogen bond with a terminal phosphate oxygen (Pautsch et al. 2013). The predicted sequence of the second putative Nile rat Gckr retrieved from Ensembl has different residues in these positions and their immediate vicinity.

### Mouse genes that appear to be absent in Nile rat

A.

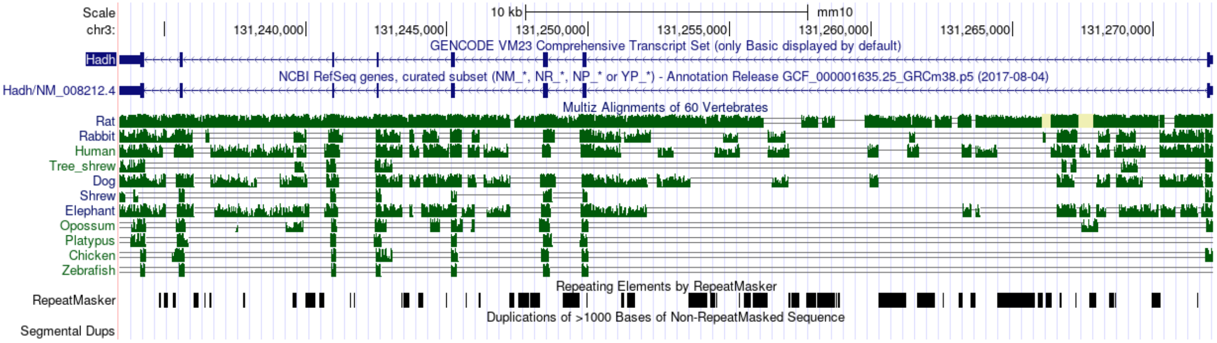

B.

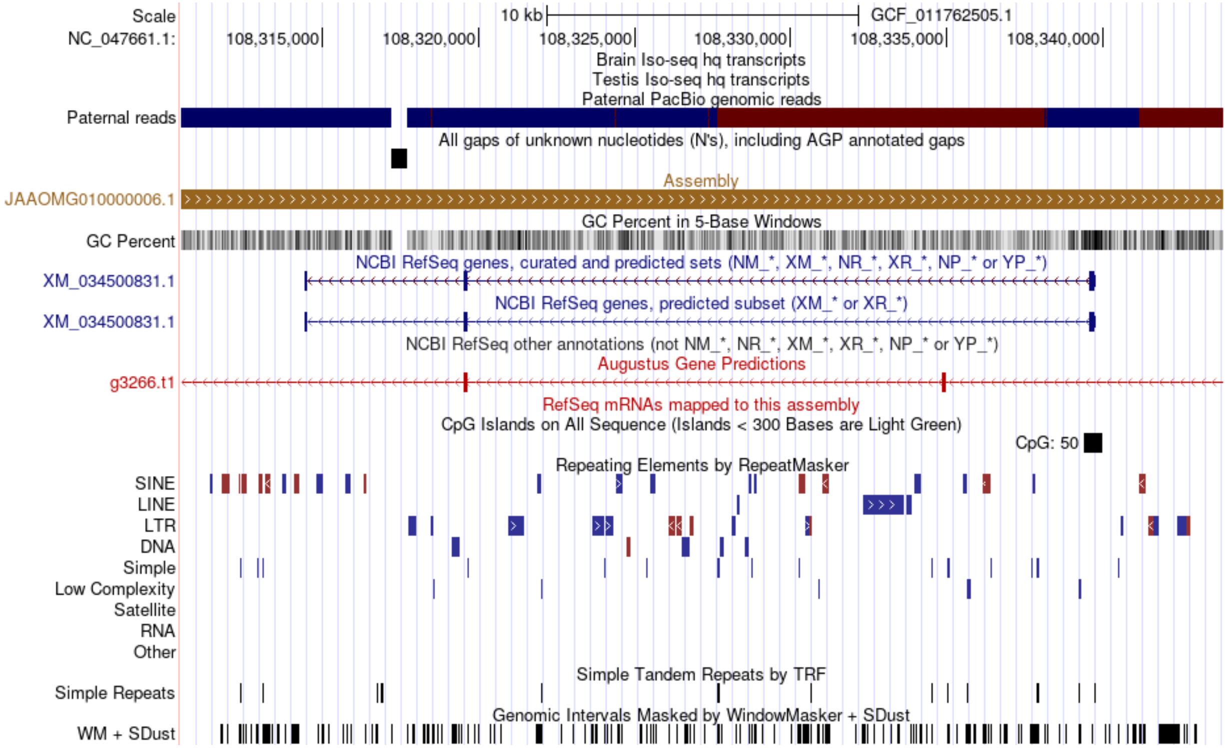


C.

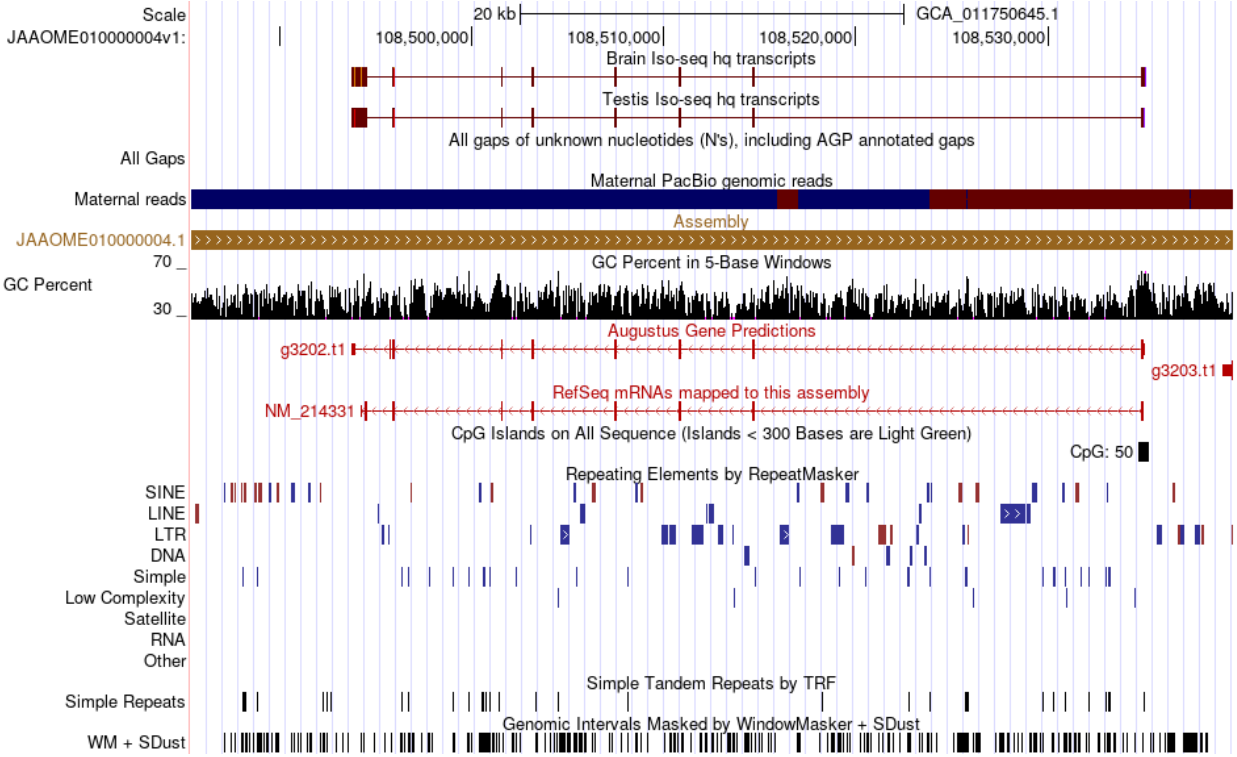


**Supplementary Figure 6. Hadh**. TOGA failed to find a functional Hadh gene in the primary assembly but did find it in the alternate assembly. **A**. Hadh gene in the mouse has 8 exons. **B**. A putative Hadh gene predicted by RefSeq in Nile rat primary haplotype assembly has only 3

exons. It has been disrupted by a gap in the assembly, which can be seen within the second intron. This gene is not supported by any Iso-seq transcripts. **C**. A full length Iso-seq transcript containing 8 exons and expressed in both brain and testis maps to the alternate haplotype assembly. This indicates haplotype variation and emphasizes the importance of analyzing both haplotypes.

A.

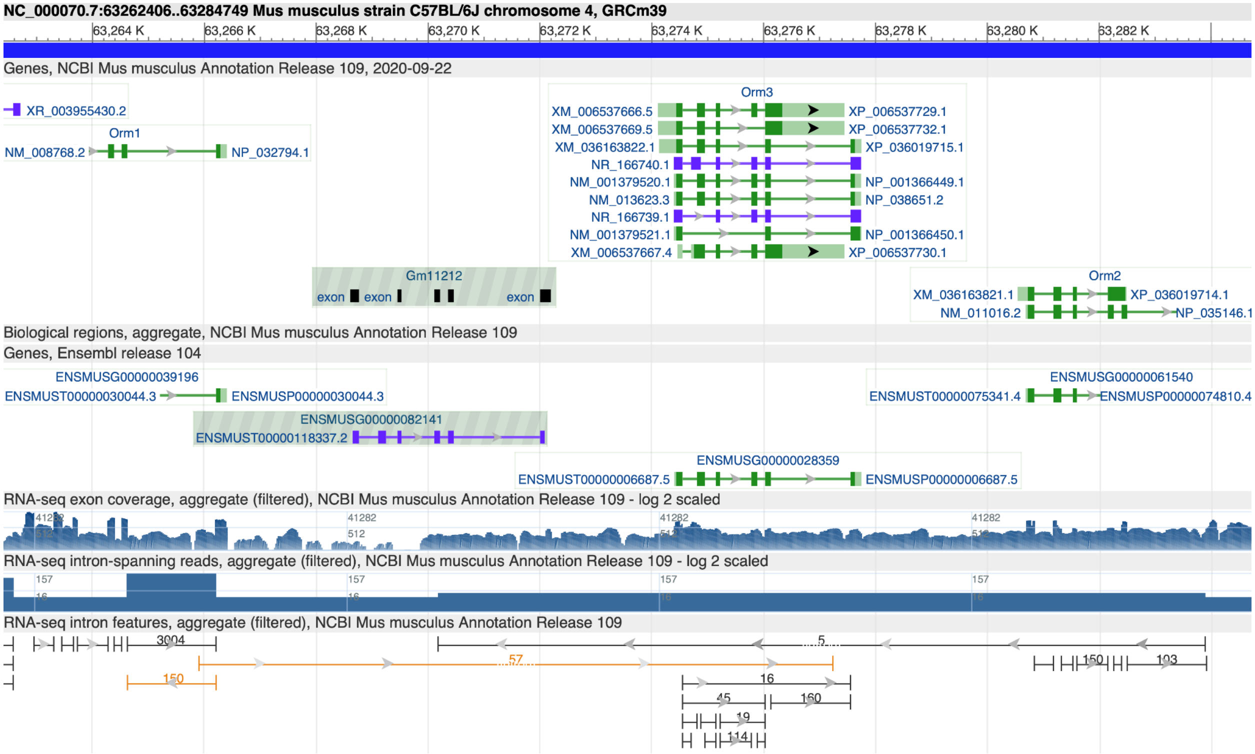


B.

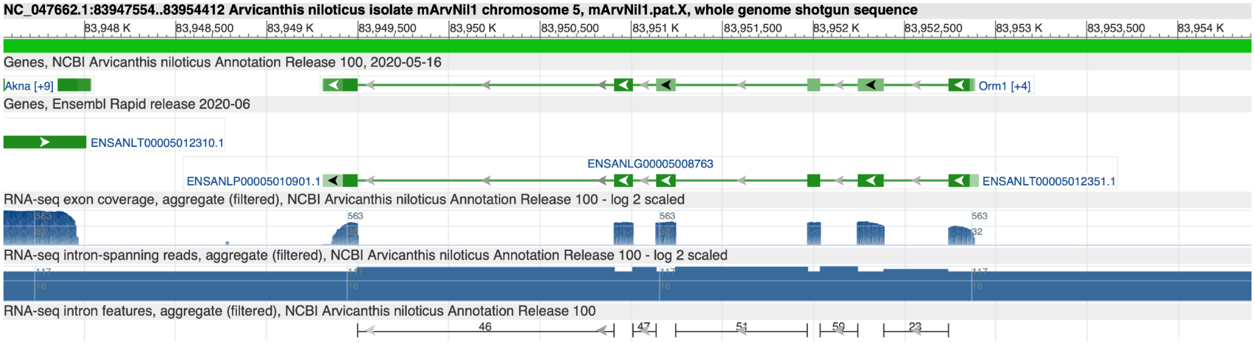

C.

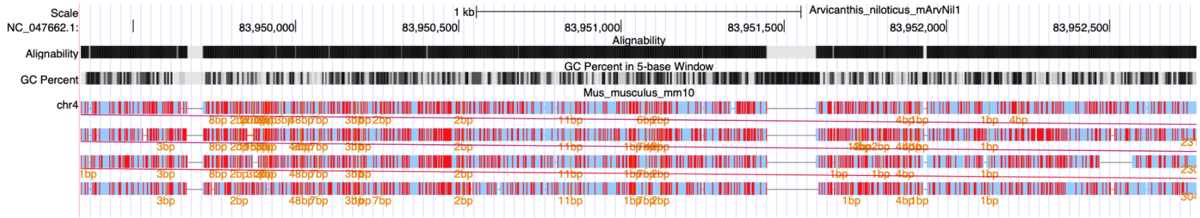


**Supplementary Figure 7. Orm2**. **A**. *Orm* gene cluster in house mouse visualized in NCBI Genome Data Viewer. The cluster contains 3 protein coding genes and 1 pseudogene. **B**. Nile rat has only one *Orm* gene. **C**. A whole genome alignment of mouse genome vs. Nile rat shows the presence of a 4-fold duplication of the *Orm* region in the mouse genome.

### GO enrichment analysis of 1601 Nile rat-specific genes

A gene set enrichment analysis of the 1601 genes that do not overlap TOGA projections from house mouse on the NaviGO server (Wei et al. 2017) revealed that several of the most highly enriched GO terms were linked to protein biosynthesis (supplementary data: https://osf.io/fh62m/). Of the top 20 statistically significant terms, the most specific protein biosynthesis-related term was GO:0022625, cytosolic large ribosomal subunit (**supplementary figure NRnotMouseTop20GOTerms**). Examination of several genes linked to this GO term revealed that these are retrogenes derived from ribosomal proteins such as L19, L21, and L23. All of these exist in multiple copies in the Nile rat genome. Multiple copies of retrogenes derived from ribosomal proteins have been known to exist in many other species of mammals (Dharia et al. 2014)

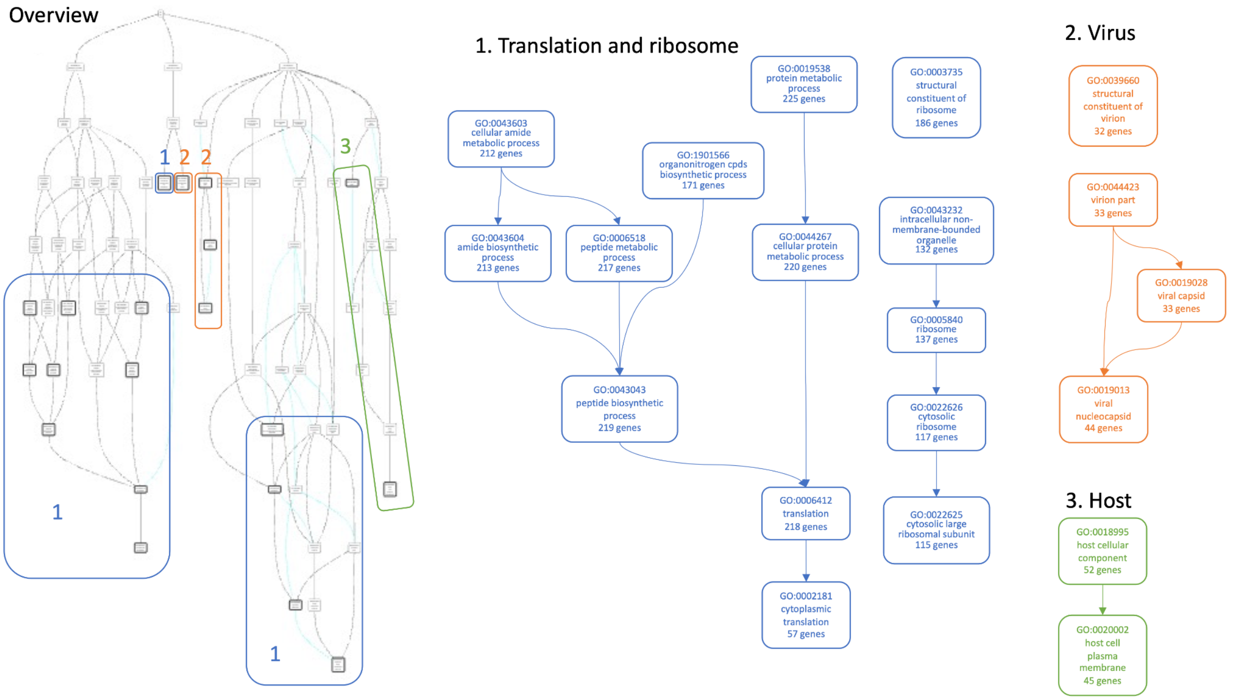

**Supplementary Figure 8. Top 20 GO terms overrepresented in Nile rat genes that do not overlap TOGA projections from the house mouse.** Overview of the 20 terms in the context of the GO hierarchy is shown on the left. The terms are grouped into 3 broad categories: 1. Translation and ribosome, 2. Virus, and 3. Host. Hierarchical relationships between terms in each category are shown on the right. The number of genes annotated with each term is shown in the corresponding box.

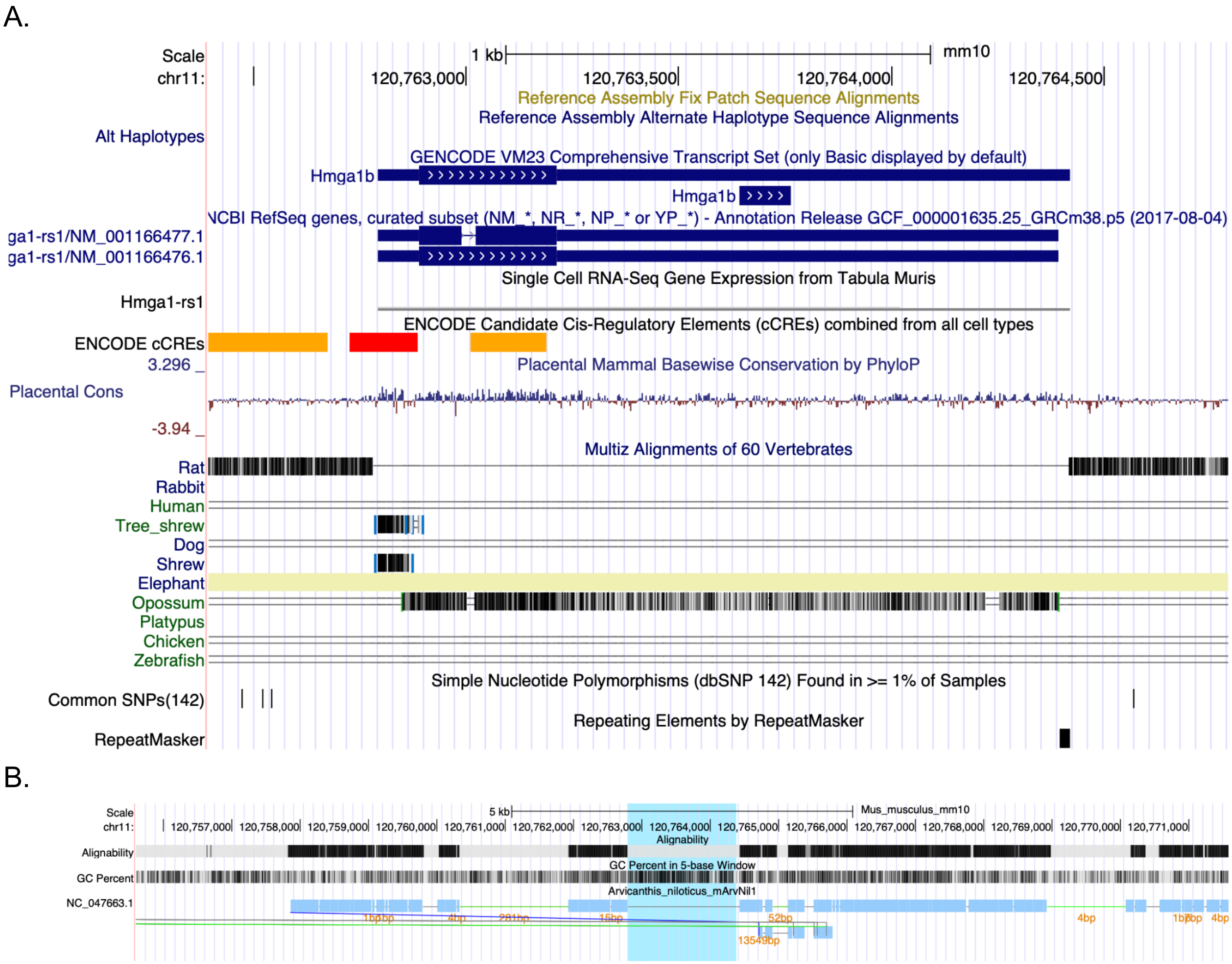


**Supplementary Figure 9. Hmga1b. A.** Mouse *Hmga1b* gene visualized in the UCSC genome browser. *Hmga1b* is a likely retrogene, as it lacks introns. Additionally, this gene is not present in the Norway rat. **B**. *Hmga1b* gene insertion in mouse relative to Nile rat. A whole genome alignment of Nile rat genome vs. mouse in the vicinity of the mouse *Hmga1b* locus (highlighted). The mouse *Hmga1b* locus does not align to the Nile rat genome, while the surrounding region aligns to chromosome 6.

A.

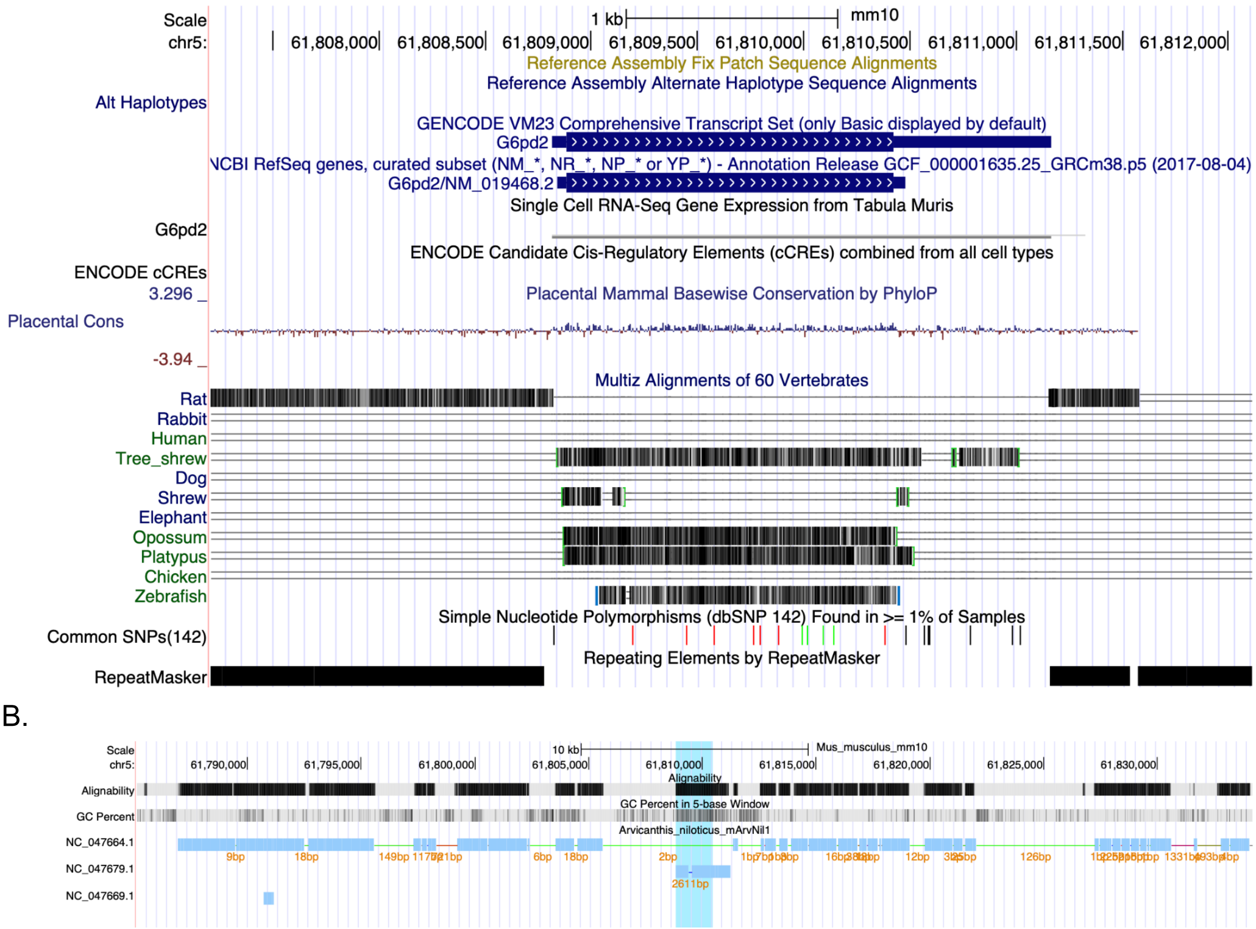


**Supplementary Figure 10. G6pd2. A.** Mouse *G6pd2* gene visualized in the UCSC genome browser. *G6pd2* is a likely retrogene, as it lacks introns. Additionally, this gene is not present in the Norway rat. **B**. *G6pd2* gene insertion in mouse relative to Nile rat. A whole genome alignment of Nile rat genome vs. mouse in the vicinity of the mouse *G6pd2* locus (highlighted). The mouse *G6pd2* locus aligns to the *G6pd* locus on Nile rat chromosome X, while the surrounding region aligns to chromosome 7. Mouse *G6pd* is likewise located on chromosome X and is the likely parent of the *G6pd2* retrogene.

### Manual modifications of haplotype assemblies for heterozygous variation detection

Having several scaffolds not anchored to chromosomes can result in overestimating overall heterozygosity and the number of structural variations. To minimize this effect, we manually modified the haplotype assemblies using a Mummer (Marçais et al. 2018) alignment. Only unanchored scaffolds longer than 500 kb, with alignment similarity over 98% and aligned blocks longer than 10 kb were used in  this analysis.  We applied the following three rules: 1) when maternal and paternal assemblies had a different orientation, the maternal assembly was reversed; 2) when small scaffolds of maternal (or paternal) assembly uniquely mapped to existing chromosomes of paternal (or maternal) and had no overlap with other mapped scaffolds, these small scaffolds were linked to existing ones with the insertion of 1000 bp gaps; 3) in more complicated cases, especially for those scaffolds that had overlaps with others, the overlapped regions were carefully trimmed as shown in (Supplemental Figure 9). This resulted in an improved version of haplotype assemblies, described by an AGP ("AGP Specification v2.1" n.d.) file available from https://osf.io/v4ypz/.

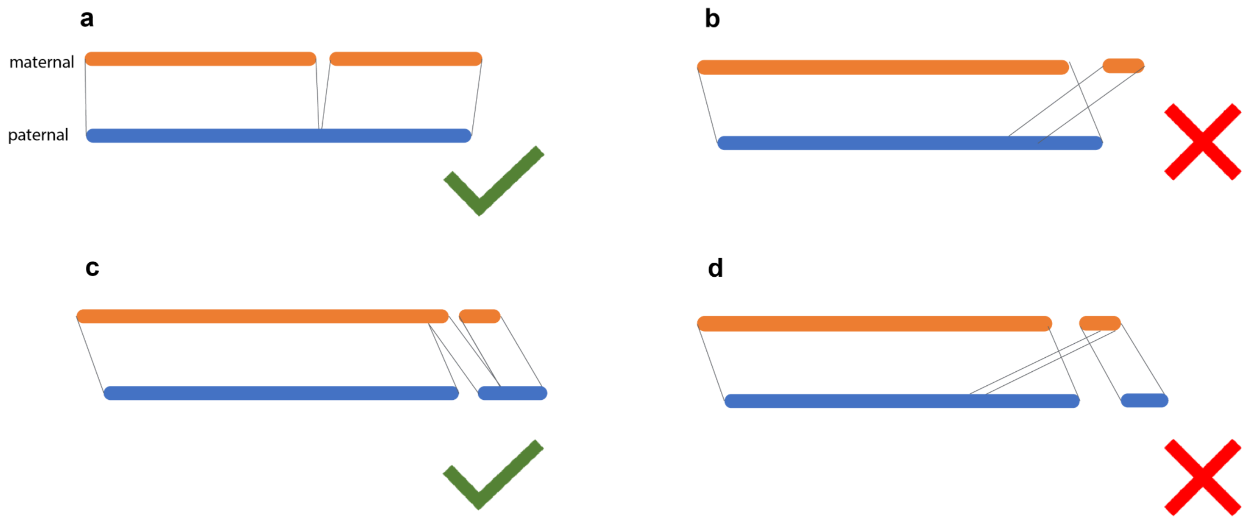


**Supplementary Figure 11. Schematic diagram of trimming alignment.** A and c can potentially link unplaced scaffolds, but b and d are not likely to be the right placements.

#### Comparison of the mouse segmental duplications computed by our workflow to prior work

To gauge how our SD annotations compare with existing ones, we compared our mm10 annotations using the sedef software with those from the UCSC genome browser generated using the software WGAC (Bailey et al. 2004). The unfiltered sedef annotations cover 149 Mb of sequence, or 5.4% of the genome, compared to 215 Mb of sequence (7.9%) by WGAC, with the majority of the difference accounted for by the Y-chromosome annotations where 69.5 Mb of additional SD are annotated by WGAC. On other chromosomes, there are roughly 20 Mb of duplicated sequences annotated by each method individually indicating that our approach mostly replicates the existing annotations of SD sequence on autosomal chromosomes.

The comparison of SD annotations for the long-read assembly of the C57BL strain of house mouse and mm10 illustrates the differences of SD resolution between the whole-genome shotgun (C57BL) assembly, and manually curated mm10 assembly. A total of 204 Mb of sequence is annotated as SD in the C57BL assembly, though the distribution of percent identity of duplications is shifted lower in C57BL (Figure 1a). Furthermore, the average length of SD annotated in C57BL is lower than mm10 (5.6 kb versus 8.4 kb), indicating that long-read sequencing assemblies may contain shorter repetitive DNA that is not removed by masking approaches and annotated as SD. When measuring gene duplication using gene multi-mapping and read depth, a similar number of duplicated genes are observed in the mm10 and C57BL (Figure 1b), indicating that the difference in total bases annotated as duplication may be accounted for by repeat masking or alignment artifacts.

#### Positively selected single amino-acid substitutions in *Xiap*

Xiap 122 G>A substitution appears to be unique among rodents. However, this residue is not well-conserved across mammalian genomes. Xiap 135T is conserved in 149 mammals, but Nile rat has 135 T>P, and four other mammals have N, I, or S. Notably, S, T, and N are all amino acids with polar uncharged side chains, but P is a nonpolar cyclic amino acid. Human variants have not been reported in residue 135, but nearby disease variants exist in positions 130, 145, and 152 (UniProt). Finally, Xiap 190Y is conserved in all mammals except African woodland thicket rat, Jamaican fruit bat, and Nile rat with Y>F substitution. The African woodland thicket rat is closely related to the Nile rat (Steppan and Schenk 2017). Although no human variant has been reported at position 190, there are nearby disease variants at positions 188 and 189 (Karczewski et al. 2020).

#### Bibliography


"AGP Specification v2.1." n.d. Accessed November 21, 2021. https://www.ncbi.nlm.nih.gov/assembly/agp/AGP_Specification/.

Bailey, Jeffrey A., Deanna M. Church, Mario Ventura, Mariano Rocchi, and Evan E. Eichler. 2004. "Analysis of Segmental Duplications and Genome Assembly in the Mouse." *Genome Research* 14 (5): 789–801.

Dharia, Asav P., Ajay Obla, Matthew D. Gajdosik, Amanda Simon, and Craig E. Nelson. 2014. "Tempo and Mode of Gene Duplication in Mammalian Ribosomal Protein Evolution." *PloS One* 9 (11): e111721.

Karczewski, Konrad J., Laurent C. Francioli, Grace Tiao, Beryl B. Cummings, Jessica Alföldi, Qingbo Wang, Ryan L. Collins, et al. 2020. "The Mutational Constraint Spectrum Quantified from Variation in 141,456 Humans." *Nature* 581 (7809): 434–43.

Marçais, Guillaume, Arthur L. Delcher, Adam M. Phillippy, Rachel Coston, Steven L. Salzberg, and Aleksey Zimin. 2018. "MUMmer4: A Fast and Versatile Genome Alignment System." *PLoS Computational Biology* 14 (1): e1005944.

Pautsch, Alexander, Nadja Stadler, Adelheid Löhle, Wolfgang Rist, Adina Berg, Lucia Glocker, Herbert Nar, et al. 2013. "Crystal Structure of Glucokinase Regulatory Protein." *Biochemistry* 52 (20): 3523–31.

Steppan, Scott J., and John J. Schenk. 2017. "Muroid Rodent Phylogenetics: 900-Species Tree Reveals Increasing Diversification Rates." *PloS One* 12 (8): e0183070.

Wei, Qing, Ishita K. Khan, Ziyun Ding, Satwica Yerneni, and Daisuke Kihara. 2017. "NaviGO: Interactive Tool for Visualization and Functional Similarity and Coherence Analysis with

Gene Ontology." *BMC Bioinformatics* 18 (1): 177.
